# Determinants of persistence in sequential effort-based decision-making

**DOI:** 10.64898/2026.05.11.723817

**Authors:** Amine Chaigneau, Riccardo Moretti, Pierpaolo Iodice, Mathias Pessiglione, Giovanni Pezzulo

## Abstract

Goal-directed behavior often requires sustained effort across a sequence of interdependent decisions, yet the determinants of persistence in such contexts remain poorly understood. Here, we investigated how individuals regulate persistence in a novel sequential effort-based task in which they controlled an avatar through successive checkpoints to reach a final goal and could make repeated attempts following failure. At each attempt, participants could choose either to persist in the same task or to disengage toward an easier but less rewarding alternative. We found that decisions to persist or disengage were jointly shaped by multiple interacting factors. Disengagement increased with task difficulty and lower skill level. It also increased with repeated attempts and varied with time-on-task, indexing fatigue, and with accumulated errors, indexing lack of progress. Conversely, proximity to the goal promoted persistence and shaped decision dynamics with greater hesitation when disengaging near the goal. Notably, clearing the first checkpoint produced a sharp increase in persistence, suggesting that early success plays a pivotal role. Furthermore, persistence reflected both retrospective and prospective evaluations of effort, with prior investment promoting commitment and anticipated effort reducing it. Finally, disengagement was preceded by short-term performance decline but not by a detectable increase in decision conflict. Together, these findings provide a comprehensive account of persistence in sequential effortful tasks, showing that decisions to persist or disengage are jointly shaped by multiple factors related to fatigue, (lack of) progress, goal proximity, and early success.

## 1 Introduction

Many goal-directed behaviors require sustained effort over time, involving a series of interdependent decisions rather than isolated choices. When pursue long-term goals, like obtaining a PhD degree, training for the Olympics or during the “hell week” of special forces, individuals must repeatedly decide whether to continue investing effort toward a goal or to disengage and redirect resources elsewhere. This raises a central question: how do individuals regulate persistence in sequential effortful tasks? Unlike one-shot decisions, persistence unfolds dynamically, as ongoing performance, internal states, and environmental conditions continuously reshape the value of persistence.

A large body of work suggests that persistence reflects a cost–benefit tradeoff, in which effort is treated as a cost that must be weighed against expected rewards (Shenhav et al., 2017; Westbrook et al., 2013; Clairis and Pessiglione, 2024; Botvinick and Cohen, 2014; Silvestrini et al., 2023; Westbrook et al., 2020; Proietti et al., 2025). For example, an influential framework suggests that the allocation of effort is governed by the expected value of control, integrating reward magnitude, effort costs, and the probability of successful goal attainment (Shenhav et al., 2013). Importantly, feedback and performance monitoring play a central role in this process, as individuals continuously evaluate ongoing outcomes to adjust effort investment (Carver and Scheier, 1990; Meyniel et al., 2013). These accounts provide a principled basis for understanding how effort is regulated, but have mainly been studied in contexts where decisions are relatively independent.

In sequential settings, however, persistence decisions depend not only on immediate costs and benefits, but also on the evolving structure of the task. As individuals progress toward a goal, multiple signals become relevant, including accumulated effort, proximity to goal completion, and recent performance outcomes. Prior work suggests that effort may accumulate over time, potentially reducing engagement (Blain et al., 2016, 2019; Matthews et al., 2023; Müller and Apps, 2019; Müller et al., 2021), while progress toward a goal can increase motivation, as formalized by goal-gradient theories (Hull, 1932; Kivetz et al., 2006; Devine and Otto, 2022). In addition, early indicators of success and ongoing performance feedback can shape expectations about future outcomes, influencing the willingness to persist (Carver and Scheier, 1990). Together, these findings suggest that persistence in sequential tasks–where individuals repeatedly decide whether to persist or disengage as effort accumulates and progress evolves–emerges from the dynamic integration of multiple effort-, progress- and feedback-related factors. How these factors interact to determine persistence decisions in sequential tasks remains not fully understood.

To address this gap, we developed a novel effort-based paradigm in which participants controlled a bird avatar navigating a sequence of obstacles within a series of trials, each corresponding to a distinct problem to solve in order to obtain a reward. Within each trial, participants progressed through successive checkpoints (CPs), which defined intermediate milestones towards the final goal, and could make repeated attempts following failure. At each attempt, they were given the opportunity to either *persist* (i.e., try again the same problem) or *disengage* (i.e., switch to an easier but less rewarding problem). We collected behavioral measures of performance and mouse-tracking indices of decision dynamics, while participants selected between persist or disengage options.

This design allows informative comparisons among effort accumulation, progress toward the goal, and performance outcomes, while acknowledging that these factors remain partially intertwined in the task structure. We aimed to assess four, not mutually exclusive hypotheses: that both repeated effort (*fatigue hypothesis*) and the accumulation of errors (*lack of progress hypothesis*) would reduce persistence, whereas both approaching the goal (*goal proximity hypothesis*) and clearing the first checkpoint within a trial (*early success hypothesis*) would increase persistence. In addition to testing these four hypotheses, we sought to address two further questions. First, we examined whether participants’ strategies varied as a function of skill level, given that individuals with different abilities may experience the task as more or less effortful. Second, we investigated the across-trial dynamics of disengagement, specifically whether the decision to disengage emerged gradually or abruptly within trials.

## 2 Methods

### 2.1 Participants

We recruited 86 participants through the Prolific platform, with eligibility criteria of native English proficiency and age between 18 and 50 years. We specified exclusion of participants who attempted to skip gameplay (near-zero engagement). Because such behavior precludes meaningful assessment of persistence, we excluded participants who completed fewer than 60 attempts overall (*≈* 2 attempts per level). The final sample consisted of 56 participants (21 men, 34 women and one who declined to state), with a median age of 35 years. All participants gave informed consent to our procedures which were approved by the Ethics Committee of the National Research Council.

### 2.2 Experimental setup

In this study, we developed a novel 2D video game using Next.js (https://nextjs.org/), inspired by Flappy Bird (Nguyen, 2013). The game required participants to control an avatar (a bird) flying through obstacles (pipes) toward a goal indicated by a coin (Figure 1A). Participants accessed the web application through their browser and used the mouse to click a jump button (red area) to make the avatar fly. The avatar moved forward automatically and was affected by gravity. If the avatar fell to the ground or hit an obstacle, the trial restarted from the last saved checkpoint (CP). Before restarting, participants were asked to choose between two options: persist in the same level or disengage by switching to a substantially easier level, while sacrificing part of the reward (5 vs. 1 coin) (Figure 1B). This design introduced multiple decision points between *persistence* and *disengagement*, depending on how participants succeeded throughout the levels.

**Fig. 1.**
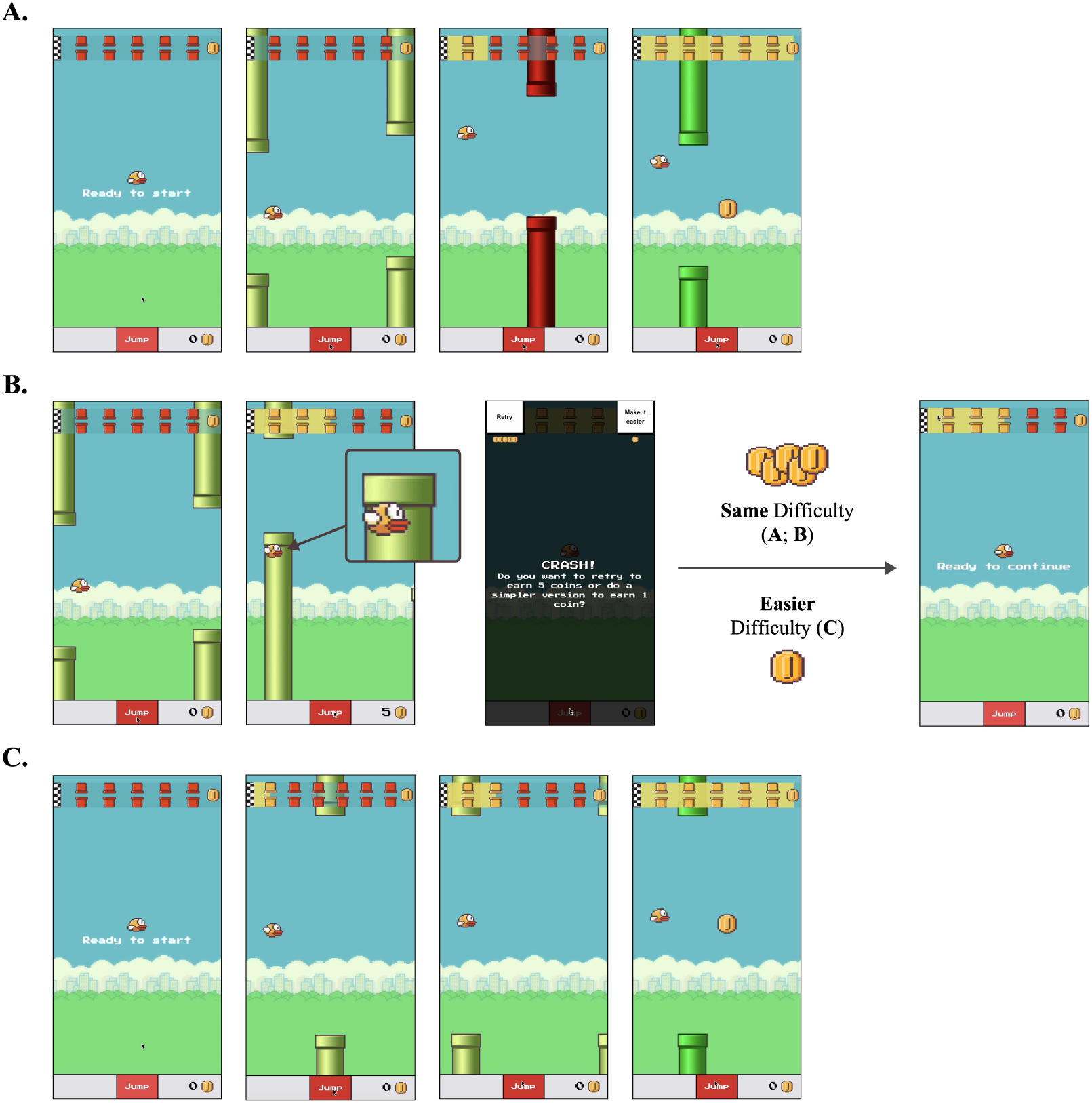
Overview of the experimental video game and task interface. (A) The main game screen. Participants controlled a bird avatar navigating through pipe obstacles toward a goal marked by a coin. The avatar moved forward automatically and was affected by gravity; participants used mouse clicks on the red jump button to maintain flight. Progress was saved at each checkpoint (CP), visually represented as red obstacles. Progress was tracked via a progress bar at the top of the game area: red obstacles indicated remaining CPs, while yellow obstacles marked previously saved CPs. The level ended when the bird reached the coin located behind the final set of light green obstacles. (B) The crash and choice interface. Following a failed attempt (e.g., collision or falling), participants immediately faced a binary choice: persist in the same level (“Retry”; potential reward of 5 coins) or disengage by switching to an easier version of the level (“Make it easier”; reward reduced to 1 coin). Regardless of choice, the level restarted from the last saved CP. (C) The easiest difficulty setting (D0), used after disengagement, which featured significantly larger horizontal and vertical gaps between obstacles.

We manipulated two independent variables: level *length* and level *difficulty*. Level length had two conditions: *short* levels with 3 CPs and *long* levels with 6 CPs. Participants were informed of the level length via an on-screen progress bar displaying CP information, including the distance to the next CP and completion status. In the game, CPs were visually represented as red obstacles, spaced by 3 obstacles between each. Difficulty was manipulated by varying the horizontal and vertical gaps between obstacles, creating two levels of difficulty. *Difficulty 1* (D1) involved larger vertical and horizontal gaps than *difficulty 2* (D2). *Easier* (D0) settings were characterized by significantly larger gaps between obstacles than both D1 and D2 (Figure 1C). The theoretical minimum duration in D0 was slightly longer than for D1 and D2 levels, so that disengaging could not be used as a shortcut to complete the experiment faster. The experimental design comprised 32 unique game levels, arranged in a 2 (Level Length: Short vs. Long) × 2 (Difficulty: D1 vs. D2) factorial design. Level order was counterbalanced across participants to control for order effects (e.g., long-first vs. short-first conditions; see Figure 2). Levels were procedurally generated within each condition; generation parameters are reported in Appendix. Before the main experiment, participants completed one level to familiarize themselves with the controls and game mechanics.

**Fig. 2.**
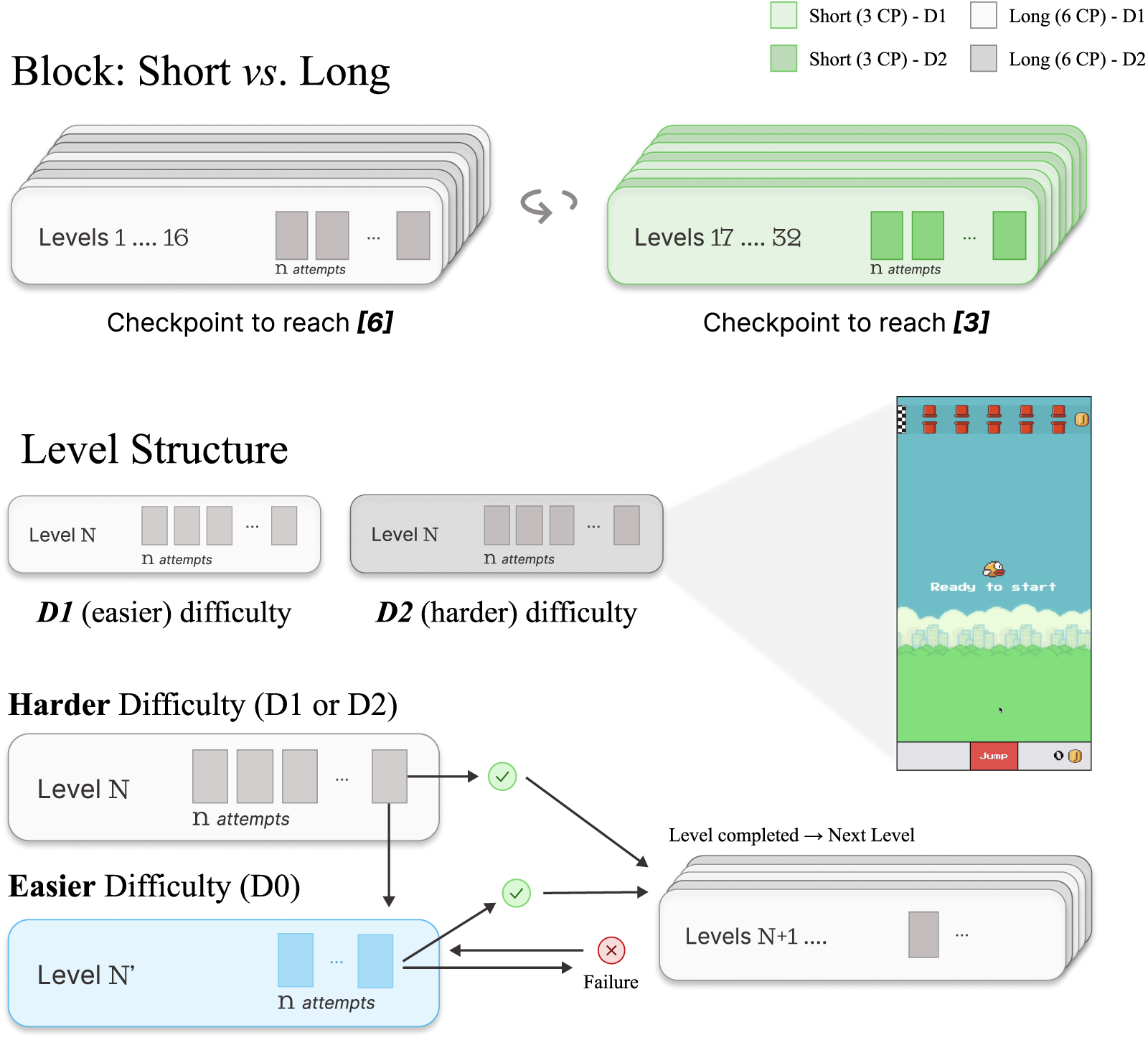
Overview of the level structure and experimental design. (Top) Participants completed 32 game levels divided into two blocks of different lengths (short vs. long), with block order counterbalanced across participants. Within each block, level difficulty (D1 or D2) was randomized across levels. (Bottom) Squares within each level card represent successive attempts, whose number varied across levels and participants. Following a crash, participants could choose either to persist in the same level or to disengage by switching to an easier version of the level (D0; see Figure 1C), in both cases resuming from the last saved checkpoint. A new level began only once the current level was fully completed, either at its original difficulty (D1 or D2) or in its easier version (D0); after a failure in D0, participants retried D0.

During the choice stage, mouse kinematics were recorded following the classic mouse-tracking framework (Freeman et al., 2010; Freeman, 2018), implemented using a *static* starting procedure (Kieslich et al., 2018). Immediately after the bird crashed, the choice interface appeared on the screen (Figure 1B). The game area window had a fixed size of 435 × 770 pixels across participants, with minimum and maximum full-screen size requirements enforced. This interface followed a static layout, ensuring a consistent starting point across trials via the red jump area (65 × 107.75 pixels) located below the gaming area. The response options, labeled *Retry* and *Make it easier*, were positioned at the top-left and top-right corners of the game area, respectively (96 × 86 pixels). For consistency in spatial analysis, the middle of the jump button was defined as the origin (0, 0), with the top-left button located at (-217.5, 835) and the top-right button at (217.5, 835). After completing the game, participants completed the Short Grit Scale (Duckworth and Quinn, 2009). The full session lasted approximately 30 minutes, and participants were paid between £6 and £12 depending on their final score (range: 32-160 points).

### 2.3 Dependent variables

#### 2.3.1 Game data

We structured the dataset at the level of individual choices, with each observation corresponding to a single attempt, defined as one instance in which a participant faced a decision point. For each attempt, we recorded the choice made (persist vs. disengage), the *attempt index* (i.e., its sequential position within the current level), the number of *obstacles passed*, and the current *CP*. CP was further transformed into *distance from goal*, defined as the number of remaining CPs at the time of choice. We also included contextual variables describing the level (i.e., length and difficulty) and the *trial index* (its sequential position within the task).

Because overall performance varied substantially across participants (*M* = 103, *SD* = 48.2; see Appendix), participants were divided into two *skill groups* based on task performance. For each participant, we computed the mean number of obstacles passed per attempt in D1 and D2 levels and performed a median split across participants. Participants above the median were classified as *skilled*, whereas those below were classified as *unskilled*. Participants in the *skilled* group achieved higher scores on average (*M* = 147, *SD* = 13.4) than those in the *unskilled* group (*M* = 59.6, *SD* = 24.2). Skill group is therefore a performance-based descriptor rather than an independent individual-difference measure. Parallel analyses treating skill as a continuous predictor (standardized mean number of obstacles passed per attempt) are reported in the Appendix. Outlier trials (top 1% of attempt counts; *>*53 attempts) were excluded from the analyses (18 trials removed).

#### 2.3.2 Mouse Tracking

Mouse cursor trajectories were recorded at a sampling rate of 10 ms throughout the choice phase. Trajectories were spatially aligned to a common starting point and direction to allow comparison across response types. Standard preprocessing procedures were applied to remove invalid trajectories and normalize movement profiles prior to analysis (Wulff et al., 2025); see the Appendix for full details. All trajectories were time-normalized to 101 equally spaced time steps using linear interpolation (Kieslich et al., 2018; Freeman et al., 2010). From each trajectory we extracted maximal absolute deviation (*MAD*) from the ideal straight-line trajectory connecting the start position to the selected response option. *MAD* provides an index of decision conflict, with larger values reflecting stronger attraction of the cursor toward the alternative response option.

### 2.4 Data analysis methods

#### 2.4.1 Modeling disengagement

Choice behavior was modeled using generalized linear mixed-effects model (GLMMs) with a logit link, fitted in *R* using the *lme4* package (Bates et al., 2015), to estimate task effects while accounting for individual variability. Continuous predictors were mean-centered and scaled to unit variance prior to analysis. Predictors reflecting fixed task structure (e.g., trial index, distance from goal) were standardized across participants, whereas performance-related predictors (e.g., attempt index, obstacles passed) were standardized within participants to isolate within-subject variability. Fixed effects were selected through nested model comparisons, retaining only predictors and interactions that significantly improved model fit based on likelihood-ratio tests (LRTs).

The final model included fixed effects for *distance from goal*, *difficulty*, and *participant skill*, as well as quadratic terms for *obstacles passed*, *attempt index*, and *trial index*. Interactions were included between *distance from goal* and *skill group*, and between *skill group* and both the quadratic effect of *attempt index* and *trial index*. The random-effects structure was selected in the same way: starting from a random intercept for participant, we tested by-participant random slopes for each within-subject predictor with LRTs. The final structure comprised a random intercept correlated with random slopes for *distance from goal* and *attempt index*, and uncorrelated random slopes for *trial index* and *obstacles passed* ; the slope for difficulty did not improve fit and was not retained.

Because *level length* and *distance from goal* are structurally collinear, they were not included simultaneously in the same model. In the primary analysis, *distance from goal* was therefore treated as a continuous predictor to capture the overall relationship between proximity to the goal and the probability of disengagement. To examine whether this global trend masked CP-specific effects, we conducted a complementary analysis in which distance was modeled as a categorical factor indexing CP position within each level-length condition. Planned contrasts compared matched checkpoints across level lengths and within levels. Auxiliary analyses testing block order effects, between-level variance, and an alternative CP-based specification of distance, together with full model-building procedures, LRTs and diagnostics, are reported in the Appendix.

#### 2.4.2 Performance dynamics preceding disengagement

To examine how performance evolved immediately prior to disengagement, we conducted a secondary analysis focusing on the three attempts preceding the attempt that ended in a decision to disengage (attempt relative to disengagement: *−*3, *−*2, *−*1). Performance (number of obstacles passed) during these attempts was modeled using a GLMM with a Poisson distribution and log link. The model included attempt relative to disengagement (entered as an ordered factor, with linear and quadratic trends), skill group, level length and all their interactions, with a random intercept for participant.

#### 2.4.3 Decision dynamics

Decision dynamics were examined using maximal absolute deviation (*MAD*) of the mouse trajectory as an index of response competition during the choice phase. Because *MAD* values were positively skewed, the measure was log-transformed prior to analysis. Log-transformed *MAD* was analyzed using linear mixed-effects models testing whether trajectory curvature varied as a function of the final decision (persist vs. disengage) and *distance from goal*, while controlling for participant *skill group*, task *difficulty*, *attempt index*, *obstacles passed*, and *trial index*. As for the choice model, the random-effects structure was selected with LRTs. It included a random intercept correlated with a by-participant random slope for decision type, and uncorrelated random slopes for distance from goal, trial index and obstacles passed. We also tested whether skill group moderated the effects of decision type and distance from goal. To examine whether decision conflict changed prior to disengagement, we conducted an additional analysis focusing on the final three attempts preceding a decision to disengage. Full model specifications and preprocessing procedures are reported in the Appendix.

## 3 Results

Participants made 9,434 decisions to persist or disengage, of which 713 (7.6%) were decisions to disengage. Because a level can end in disengagement only once, this corresponds to 40.5% of levels on average being completed in their easier version, with large differences between participants (range: 0–100%; skilled: *M* = 20.3%, *SD* = 25.7; unskilled: *M* = 60.7%, *SD* = 36.3). Disengagement was more frequent in harder than in easier levels, and in long than in short levels (D2: 53.5% of long and 46.5% of short levels; D1: 36.2% and 27.0%, respectively).

### 3.1 Characterizing the skill groups

Participants were divided into two skill groups by a median split on the mean number of obstacles passed per attempt. Because the groups are defined by performance, differences in progression between them are expected by construction. We report them here to describe how the two groups experienced the task, not as independent findings (Figure 3). Skilled participants were more likely than unskilled participants to reach later CPs at least once, and this difference increased with CP depth, particularly in long levels (Figure 3A). We also examined within-attempt progression as a function of the CP from which an attempt began. From the same starting CP, skilled participants tended to progress farther into the level than unskilled participants (Figure 3D). Transition matrices (Figure 3E-F) illustrate this pattern descriptively: skilled participants showed a higher probability of transitioning to more distant CPs, whereas unskilled participants more often advanced by a single CP before failing. To quantify this difference, we modeled CP progression within attempts using a mixed-effects model predicting the number of CPs gained during an attempt as a function of skill group, starting CP, level length, and difficulty, with a random intercept for participant. As expected, skilled participants advanced significantly farther per attempt than unskilled participants (see the Appendix).

**Fig. 3.**
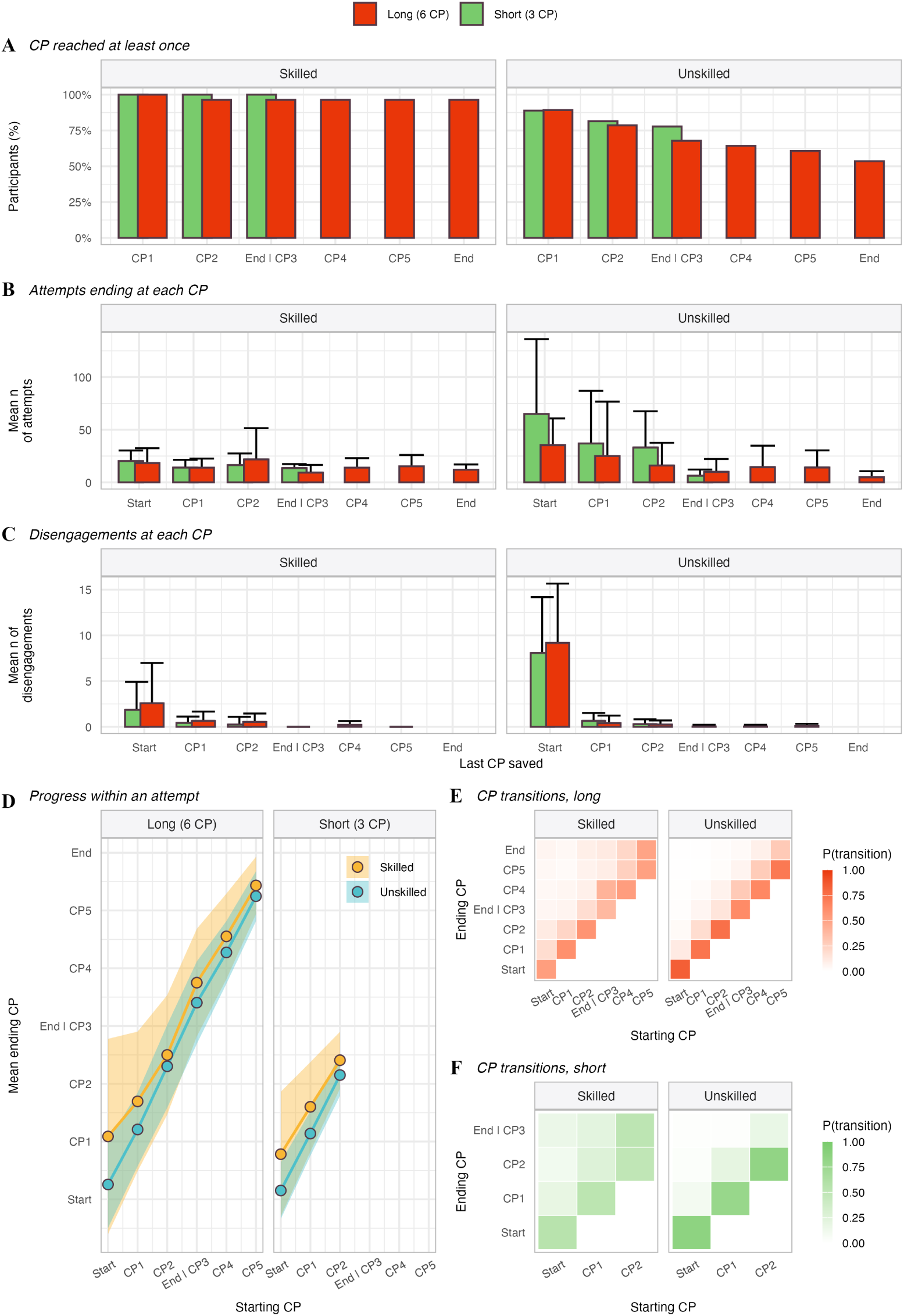
Progression dynamics and attempt outcomes as a function of skill group and level length. Because skill groups were defined by performance, differences between them are expected by construction. (A) The proportion of participants who reached each checkpoint (CP) at least once, that is, whose furthest CP across all levels of a given length was at or beyond it. (B) The mean number of attempts ending at each CP, due to either a crash, a disengagement, or successful completion. (C) The mean number of disengagements at each CP, indexed by the last CP saved. In (B) and (C), counts were computed per participant and averaged within each skill group and level length; error bars indicate one standard deviation across participants. (D) The average progression within an attempt as a function of the starting CP, computed across all attempts; shaded areas represent one standard deviation. The ending CP is the furthest CP reached within an attempt, which equals the starting CP when no progress was made. (E, F) Transition probabilities between CPs in long (E) and short (F) levels, computed per participant and normalized within each starting CP, then averaged across participants within each skill group. Note that CP3 in long levels corresponds to the end of short levels.

### 3.2 Disengagement increases with task difficulty and lower skill

Participants in the *unskilled* group were more likely to disengage than those in the *skilled* group (Unskilled: *β* = 2.41, *p < .*001). In addition, easier trials were associated with a lower probability of disengagement (Difficulty D1: *β* = *−*0.34, *p < .*01), indicating that disengagement increased with task difficulty.

The same results were obtained when skill was treated as a continuous predictor rather than a group split (see Appendix). Both effects are expected, since lower skill and higher difficulty make failure more likely and therefore raise the expected cost of completing a level. Consistent with this, replacing skill and difficulty with a single estimate of the likelihood of failing at each obstacle did not reduce model fit (*χ*^2^(2) = 1.11, *p* = .575; see Appendix). Self-reported grit was not associated with participants’ overall disengagement rate (*ρ* = .06, *p* = .699) or with skill (*ρ* = *−.*02, *p* = .906; see Appendix). We next examined whether persistence dynamics were additionally shaped by repeated engagement and accumulated effort.

**Table 1.**
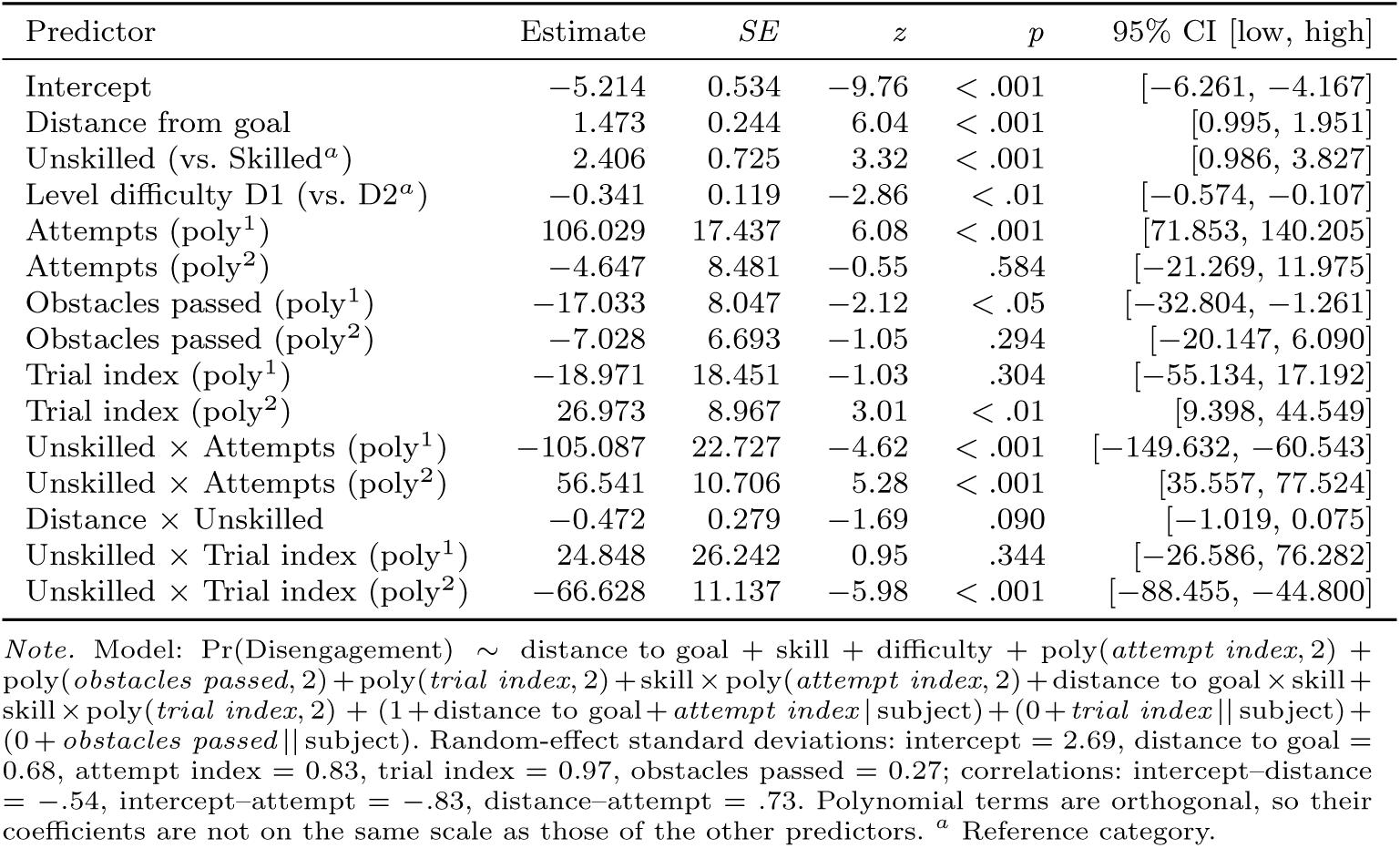
Fixed-effect estimates for the final GLMM predicting disengagement behavior (logit link). Estimates and 95% Wald CIs are on the log-odds scale.

| Predictor | Estimate | SE | z | p | 95% CI [low, high] |
| --- | --- | --- | --- | --- | --- |
| Intercept | -5.214 | 0.534 | -9.76 | < .001 | [-6.261, -4.167] |
| Distance from goal | 1.473 | 0.244 | 6.04 | < .001 | [0.995, 1.951] |
| Unskilled (vs. Skilled<sup>a</sup>) | 2.406 | 0.725 | 3.32 | < .001 | [0.986, 3.827] |
| Level difficulty D1 (vs. D2<sup>a</sup>) | -0.341 | 0.119 | -2.86 | < .01 | [-0.574, -0.107] |
| Attempts (poly<sup>1</sup>) | 106.029 | 17.437 | 6.08 | < .001 | [71.853, 140.205] |
| Attempts (poly<sup>2</sup>) | -4.647 | 8.481 | -0.55 | .584 | [-21.269, 11.975] |
| Obstacles passed (poly<sup>1</sup>) | -17.033 | 8.047 | -2.12 | < .05 | [-32.804, -1.261] |
| Obstacles passed (poly<sup>2</sup>) | -7.028 | 6.693 | -1.05 | .294 | [-20.147, 6.090] |
| Trial index (poly<sup>1</sup>) | -18.971 | 18.451 | -1.03 | .304 | [-55.134, 17.192] |
| Trial index (poly<sup>2</sup>) | 26.973 | 8.967 | 3.01 | < .01 | [9.398, 44.549] |
| Unskilled × Attempts (poly<sup>1</sup>) | -105.087 | 22.727 | -4.62 | < .001 | [-149.632, -60.543] |
| Unskilled × Attempts (poly<sup>2</sup>) | 56.541 | 10.706 | 5.28 | < .001 | [35.557, 77.524] |
| Distance × Unskilled | -0.472 | 0.279 | -1.69 | .090 | [-1.019, 0.075] |
| Unskilled × Trial index (poly<sup>1</sup>) | 24.848 | 26.242 | 0.95 | .344 | [-26.586, 76.282] |
| Unskilled × Trial index (poly<sup>2</sup>) | -66.628 | 11.137 | -5.98 | < .001 | [-88.455, -44.800] |

*Note.* Model: $\text{Pr}(\text{Disengagement}) \sim \text{distance to goal} + \text{skill} + \text{difficulty} + \text{poly}(\text{attempt index}, 2) + \text{poly}(\text{obstacles passed}, 2) + \text{poly}(\text{trial index}, 2) + \text{skill} \times \text{poly}(\text{attempt index}, 2) + \text{distance to goal} \times \text{skill} + \text{skill} \times \text{poly}(\text{trial index}, 2) + (1 + \text{distance to goal} + \text{attempt index} | \text{subject}) + (0 + \text{trial index} | \text{subject}) + (0 + \text{obstacles passed} | \text{subject})$ . Random-effect standard deviations: intercept = 2.69, distance to goal = 0.68, attempt index = 0.83, trial index = 0.97, obstacles passed = 0.27; correlations: intercept-distance = $-.54$ , intercept-attempt = $-.83$ , distance-attempt = $.73$ . Polynomial terms are orthogonal, so their coefficients are not on the same scale as those of the other predictors. <sup>a</sup> Reference category.

### 3.3 Repeated attempts increase disengagement probability

Within trials, the probability of disengagement increased with *attempt index*, with a profile that differed by skill group. Among skilled participants, disengagement increased linearly across attempts (Attempts (poly^1^): *β* = 106.03, *p < .*001), with no quadratic component (Attempts (poly^2^): *β* = *−*4.65, *p* = .584). Among unskilled participants, this linear increase was attenuated (Unskilled *×* Attempts (poly^1^): *β* = *−*105.09, *p < .*001) and a quadratic component emerged (Unskilled *×* Attempts (poly^2^): *β* = 56.54, *p < .*001). Thus, unskilled participants were initially more likely to disengage early, became more persistent once engaged, and then showed renewed disengagement after repeated failures (Figure 4F).

**Fig. 4.**
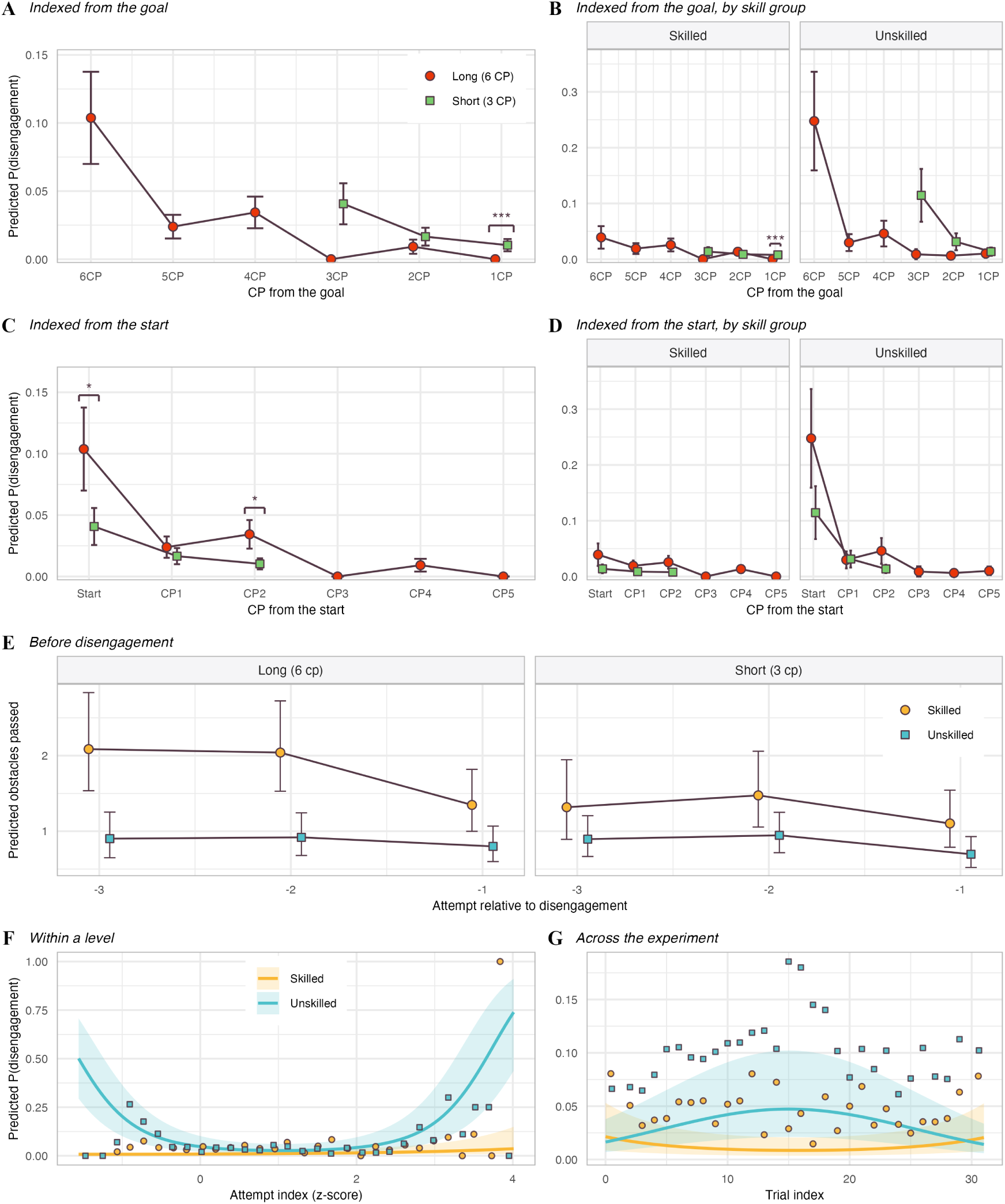
Disengagement as a function of task structure, recent performance and time. (A, B) Predicted probability of disengagement with CPs indexed from the goal (distance remaining from the goal), for all participants (A) and by skill group (B). (C, D) The same data indexed from the start of the level (distance progressed from the start), for all participants (C) and by skill group (D). Each point represents the model-predicted probability derived from the binomial GLMM, with error bars indicating standard errors. Asterisks denote significant pairwise contrasts between short and long levels at matched checkpoints (Holm-corrected): * *p < .*05, *** *p < .*001; contrasts at 1CP rest on few disengagements. (E) The predicted number of obstacles passed in the three attempts preceding a disengagement. Points represent model-estimated marginal means from the Poisson GLMM, and error bars indicate 95% confidence intervals. (F) Predicted probability of disengagement across attempts within a level (standardized attempt index), modeled with a quadratic term. (G) Predicted probability of disengagement across trials. In (F) and (G), predictions are from the main model; solid lines and shaded ribbons represent model predictions and 95% confidence intervals, and dots show observed mean probabilities at binned values, providing an empirical check of model fit.

### 3.4 Goal proximity reduces disengagement

Our analysis also reveals that *distance from goal* positively predicted the log-odds of disengagement (distance from goal: *β* = 1.47, *p < .*001), indicating that participants were less likely to disengage when they were closer to the goal. This effect was not significantly moderated by skill group (distance from goal *×* Unskilled: *β* = *−*0.47, *p* = .090).

This pattern was also reflected in the negative linear effect of *obstacles passed* within an attempt (Obstacles passed (poly^1^): *β* = *−*17.03, *p < .*05), indicating that greater progress in one attempt was associated with reduced probability of disengagement.

### 3.5 Clearing the first checkpoint increases persistence

To test whether early success exerted an effect beyond overall goal distance, we examined CP-specific contrasts within short and long levels. A clear temporal pattern emerged: disengagement was highest before the first CP was cleared and dropped sharply once the first CP had been passed (Figure 4A; Table 2). This early drop in disengagement probability was significant in both long levels (Long 6CP vs. 5CP: *z* = 6.23, *p_holm_ < .*001) and short levels (Short 3CP vs. 2CP: *z* = 3.87, *p_holm_ < .*001). In long levels, a further decrease was observed between the final two checkpoints (Long 2CP vs. 1CP: *z* = 4.79, *p_holm_ < .*001), although this contrast rests on very few disengagement events. No corresponding decrease was found in short levels (Short 2CP vs. 1CP: *z* = 1.37, *p_holm_* = .172).

**Table 2.** Significant planned contrasts comparing disengagement probabilities between short and long levels and across checkpoints.

| Contrast | Odds ratio | SE | $z$ | $p_{holm}$ |
| --- | --- | --- | --- | --- |
| *Distance from goal* |  |  |  |  |
| Short vs Long (1CP) | 2719.12 | 4265.21 | 5.04 | < .001 |
| Long 6CP vs 5CP | 4.73 | 1.179 | 6.23 | < .001 |
| Long 2CP vs 1CP | 2421.60 | 3937.43 | 4.79 | < .001 |
| Short 3CP vs 2CP | 2.52 | 0.600 | 3.87 | < .001 |
| *Distance from start* |  |  |  |  |
| Short vs Long (Start, CP0) | 0.37 | 0.132 | -2.79 | < .05 |
| Short vs Long (CP2) | 0.29 | 0.142 | -2.53 | < .05 |
| *Skill-specific* |  |  |  |  |
| Short vs Long (1CP) — Skilled | 5.52e+06 | 1.66e+07 | 5.17 | < .001 |

*Note.* Model: $\text{Pr}(\text{Disengagement}) \sim \text{distance by condition} + \text{distance by condition} \times \text{skill} + (1 + \text{distance to goal} + \text{level length} | \text{participant})$ . Odds ratios are reported as exponentiated pairwise contrasts from the underlying logit model. $z$ statistics and $p$ values are based on the corresponding contrasts on the log-odds scale. Very large or very small odds ratios reflect sparse disengaging events in some cells and should be interpreted with caution. For clarity, only statistically significant contrasts are shown; the complete set of contrasts is reported in Appendix. $p$ values are Holm-corrected.

In contrast, intermediate checkpoint comparisons within long levels were not significant, indicating that the transition in disengagement dynamics occurred abruptly early in the level. By comparison, the decline in disengagement across checkpoints in short levels was more gradual. Together, these results indicate a pronounced effect of clearing the first checkpoint on persistence, above and beyond the global influence of goal proximity. This drop is unlikely to reflect sensorimotor learning alone. Although performance improved over the experiment (*p < .*001), the drop was not smaller when a level length was encountered in the second block than in the first (checkpoint *×* block position: long, *p* = .620; short, *p* = .798; see Appendix).

Taken together, these results indicate that disengagement decisions are jointly shaped by effort costs, fatigue dynamics, and progress-related signals, including both proximity to the goal and first checkpoint within a level.

### 3.6 Retrospective and prospective effort jointly shape persistence

We further examined whether disengagement decisions depended not only on absolute distance from goal but also on the broader trial context. Specifically, the effort already invested (retrospective context) and the effort remaining to reach the goal (prospective context). Because identical checkpoint distances can occur in both short and long trials, our design allows comparisons of states with the same distance from goal but different effort histories. For example, being 3 CPs from the goal can occur either early in a short trial or after several checkpoints have already been cleared in a long trial.

When comparing identical distances from the goal across trial lengths, participants were less likely to disengage in *Long* trials than in *Short* trials when one CP remained (Short vs. Long at 1CP: *z* = 5.04, *p_holm_ < .*001; Figure 4A). This difference was not significant at the other matched distances (2CP: *p_holm_* = .350; 3CP: *p_holm_* = .075). These results suggest that persistence decisions depend partly on the effort already invested in the trial.

When checkpoints were indexed from the start of the level, the opposite pattern emerged. Participants were more likely to disengage in *Long* trials than in *Short* trials at equivalent progress positions (Start/CP0: *z* = *−*2.79, *p_holm_ < .*05; CP2: *z* = *−*2.53, *p_holm_ < .*05; Figure 4C).

### 3.7 Performance decline precedes disengagement

We next tested whether the decision to disengage was preceded by a decline in performance. We found a clear pre-disengagement pattern: performance declined as participants approached the decision of switching to easier settings. The number of obstacles passed decreased significantly across the final three attempts (*β* = *−*0.31, *p < .*01, Figure 4E), indicating that disengagement was typically preceded by a short sequence of poorer outcomes. As expected from the main analyses, unskilled participants also showed lower overall performance than skilled participants (*β* = *−*0.72, *p < .*001), and performance was lower in short than in long levels (*β* = *−*0.33, *p < .*01).

Critically, the decline in performance preceding disengagement did not differ reliably as a function of skill group or level length (all interactions with attempt relative to disengagement, *p*s *> .*17). Thus, although disengagement was typically preceded by performance deterioration, the immediate trajectory leading up to disengagement was broadly similar across groups and conditions.

This pre-disengagement decline is consistent with the main decision model, in which the number of obstacles passed negatively predicted disengagement, indicating that poorer performance was associated with a higher probability of disengagement.

### 3.8 Disengagement shows a mid-task peak with time-on-task

At the between-trial level, *trial index* showed a significant quadratic effect (Trial index (poly^2^): *β* = 26.97, *p < .*01), indicating that disengagement changed non-linearly over the course of the experiment, consistent with fatigue accumulation during the task. This temporal profile differed across skill groups. Although the linear interaction between *trial index* and skill group was not significant (Unskilled *×* Trial index (poly^1^): *β* = 24.85, *p* = .344), the quadratic interaction was negative and significant (Unskilled *×* Trial index (poly^2^): *β* = *−*66.63, *p < .*001), indicating an inverted-U pattern among unskilled participants, with the probability of disengagement peaking around the middle of the experiment (Figure 4G). We consider possible accounts of this pattern in the Discussion.

### 3.9 Decision dynamics reveal conflict during persistence and disengagement

To examine the dynamics of decision formation, we analyzed cursor trajectories during the choice phase. Trajectories were significantly modulated by both the final decision and distance from goal (Table 3). Overall, persistence decisions produced significantly more direct cursor trajectories than disengagement decisions (*β* = *−*0.88, *p < .*001). In addition, the effect of distance from goal depended on the decision (decision *×* distance: *β* = 0.09, *p < .*01; Figure 5). Persistence decisions showed direct trajectories at all distances from the goal (*β* = 0.01, *p* = .554). In contrast, disengagement decisions exhibited greater curvature when made near the goal (*β* = *−*0.09, *p < .*05), suggesting increased hesitation when disengaging despite being close to success. Together, these results indicate that goal proximity not only influenced the likelihood of persisting but also shaped the dynamics of the decision process itself. Skill group did not influence trajectory curvature, either alone or in interaction with decision type or distance from goal (all *p*s *> .*24; see Appendix).

**Fig. 5.**
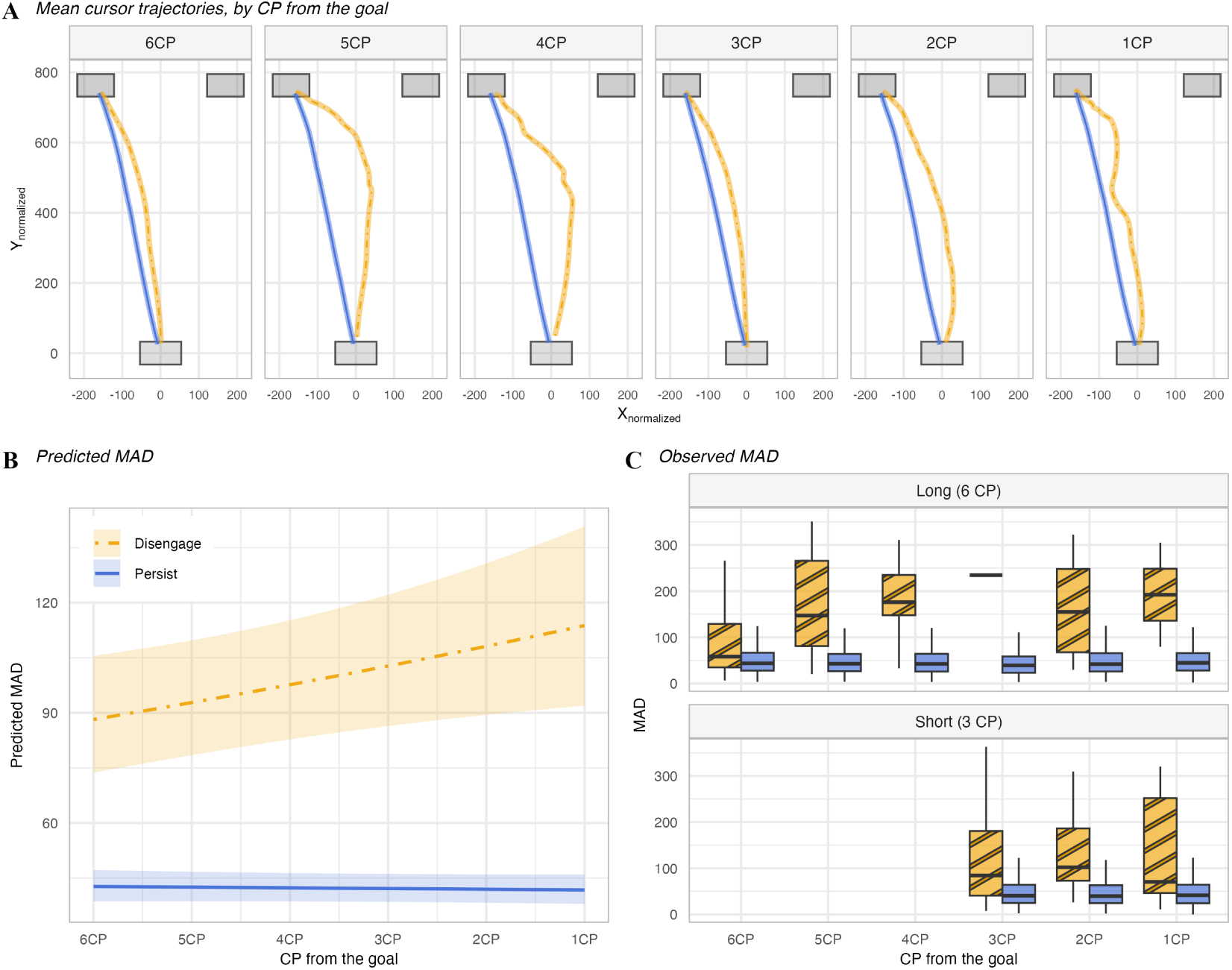
Decision dynamics during persistence and disengagement. (A) Average cursor trajectories during the choice phase as a function of distance from the goal. All trajectories were spatially realigned to a common starting point, time normalized and flipped to a common side to enable comparison across trials. (B) Model-predicted maximal absolute deviation (*MAD*) from the ideal straight-line trajectory as a function of distance to the goal, back-transformed from the log scale. Higher *MAD* values indicate stronger attraction toward the alternative response option and therefore greater decision conflict. Shaded areas represent 95% confidence intervals. (C) The distribution of observed *MAD* values across checkpoints for each decision and level length; boxplots summarize all valid trajectories.

**Table 3.** Fixed-effect estimates for the linear mixed-effects model predicting trajectory curvature (log-transformed MAD).

| Predictor | Estimate | SE | *t* | *p* |
| --- | --- | --- | --- | --- |
| Intercept | 4.653 | 0.101 | 46.27 | < .001 |
| Persist (vs. Disengage<sup>a</sup>) | −0.882 | 0.097 | −9.10 | < .001 |
| Distance from goal | −0.085 | 0.035 | −2.43 | < .05 |
| Unskilled (vs. Skilled<sup>a</sup>) | −0.079 | 0.083 | −0.96 | .344 |
| Level difficulty D1 (vs. D2<sup>a</sup>) | 0.021 | 0.019 | 1.12 | .265 |
| Attempt index | −0.026 | 0.009 | −2.79 | < .01 |
| Trial index | −0.049 | 0.019 | −2.56 | < .05 |
| Obstacles passed | 0.037 | 0.011 | 3.46 | < .01 |
| Persist × Distance from goal | 0.093 | 0.035 | 2.61 | < .01 |

*Note.* The model included a random intercept correlated with a by-participant random slope for decision type, and uncorrelated random slopes for distance from goal, trial index and obstacles passed. <sup>a</sup> Reference category.

We next examined whether decision conflict changed across the final attempts preceding a disengagement decision by modeling trajectory curvature across the last three attempts before disengagement. No significant effects of attempt relative to disengagement were observed (*p >* .26), indicating that trajectory curvature did not systematically change prior to disengagement. Full model specifications and results for the mouse-tracking analyses are reported in the Appendix.

## 4 Discussion

To investigate how individuals regulate persistence during sequential effortful tasks, we developed a novel effort-based paradigm in which participants controlled a bird avatar navigating a series of obstacles to obtain a reward at the end of each trial. Participants could disengage at any point and switch to an easier, less rewarding version of the task, allowing us to examine how persistence in goal-directed behavior evolves over time.

Our goal was to test four (not mutually exclusive) hypotheses about the regulation of persistence: that repeated effort (*fatigue hypothesis*) and accumulating errors (*lack of progress hypothesis*) would reduce engagement, and that both approaching the goal (*goal proximity hypothesis*) and clearing the first checkpoint within a trial (*early success hypothesis*) would increase persistence. In addition, our design allowed us to examine how persistence decisions are shaped by both *retrospective effort* (effort already invested) and *prospective effort* (effort remaining to reach the goal), and to characterize the dynamics of these decisions through mouse-tracking measures of response competition.

Overall, our results indicate that persistence decisions emerge from the interaction of multiple effort- and progress-related factors – related to fatigue, lack of progress, goal proximity, and clearance of the first checkpoint – which also shape the dynamics of the decision process itself.

Our results showed that the probability of disengagement increased with repeated attempts within trials and varied with progression through the experiment. These time-dependent effects suggest that sustained engagement in the task progressively altered participants’ willingness to persist. Because both attempt number and trial index capture the repeated exertion of effort, these variables may reflect the accumulation of fatigue over time, a process known to influence effort-based decision making (Blain et al., 2016, 2019; Matthews et al., 2023; Shenhav et al., 2017; Iodice et al., 2017). Specifically, fatigue may increase the subjective cost of exerting effort, thereby reducing the net value of pursuing the current goal (i.e., goal value minus effort cost), in line with the *fatigue hypothesis* and recent theoretical accounts of effort-based decision making (Pessiglione et al., 2025).

Among unskilled participants, disengagement also followed an inverted-U profile across the experiment, peaking around its midpoint. Several non-exclusive accounts could explain this pattern. A transient fatigue account holds that accumulated effort raised the subjective cost of persisting in the middle of the task; however, fatigue alone predicts a monotonic increase in disengagement, and would require an additional end-of-task effect to explain its later decline. The pattern also mirrors the “stuck-in-the-middle” effect, whereby motivation is highest near the start and the end of goal pursuit, when progress is most salient relative to the closest reference point, and lowest in between (Bonezzi et al., 2011; Pezzulo et al., 2026; Pisauro et al., 2026). Relatedly, opportunity-cost accounts attribute increased effort near the end of a task to changes in the cost of the time it requires (Emanuel et al., 2022), which would explain the late decline in disengagement without assuming recovery from fatigue. These accounts could be distinguished by manipulating session length or the information participants receive about it. Finally, because performance improved over the experiment and level length changed between the two halves of the session, learning and the block transition may also have contributed to this pattern.

Our results also showed that the probability of disengagement was significantly modulated by factors related to difficulty, performance monitoring and accumulating errors, above and beyond fatigue and time-on-task. Specifically, participants were more likely to disengage in more difficult levels, when their skill was lower and when they made repeated errors. These observations are in keeping with the *lack of progress hypothesis*, suggesting that the (perceived) low likelihood of making progress within a challenging task is a significant determinant of the decision to disengage from it. They are also in line with motivation intensity theory (Brehm and Self, 1989), which holds that effort increases with task difficulty only as long as success is possible and worthwhile, and is withdrawn otherwise. Extensions of this theory propose that lower ability and fatigue both increase the subjective difficulty of a task (Wright, 2008), which may explain why disengagement was most frequent among less skilled participants, in harder levels, and after repeated failures.

Beyond effort accumulation and lack of progress, persistence decisions were strongly shaped by progress signals within the task. We observed that the probability of disengaging decreased as participants advanced within an attempt, with greater numbers of successfully passed obstacles associated with a lower likelihood of disengaging. Because within-attempt progress directly reflects the effectiveness of ongoing control, this pattern suggests that participants were more willing to invest effort when their actions produced reliable forward progress, consistent with accounts of *control efficacy* guiding effort allocation (Shenhav et al., 2021b).

A similar effect emerged at the level of overall goal proximity. Participants were less likely to disengage when they were closer to the goal, indicating that proximity to reward counteracted disengagement. Together, these results indicate that persistence is not determined solely by accumulated effort costs, but is continuously modulated by signals of progress toward goal completion, in line with the *goal proximity hypothesis*.

Importantly, progress may also act as a proxy for expected success probability. Each successfully passed obstacle provides evidence about current control efficacy, allowing participants to update their estimated probability of successful completion. Thus, persistence may reflect not only goal proximity, but also belief updating about future outcomes, whereby increased confidence in success promotes continued engagement (Shenhav et al., 2013, 2021a; Pezzulo et al., 2024, 2018; Rigoli and Pezzulo, 2022).

Goal proximity also shaped the dynamics of the decision process itself. Cursor trajectories revealed that persistence decisions were direct at all distances from the goal, indicating low decisional conflict (Freeman, 2018). In contrast, disengagement decisions made close to the goal showed increased trajectory curvature, suggesting greater hesitation when abandoning a nearly completed goal. These findings indicate that progress signals modulate not only choice outcomes but also the degree of competition between response alternatives during decision formation. Specifically, this competition may arise because disengagement decisions made near the goal reflect a conflict between a locally high value of disengaging (e.g., following recent failures) and a simultaneously high expected value of continuing, given the proximity to achieve the goal. More broadly, these results are consistent with *goal-gradient* accounts of motivation, whereby the value of goal-directed actions increases as individuals approach reward (Hull, 1932; Kivetz et al., 2006). Our findings extend this literature to persistence decisions by showing that such effects are expressed not only at the level of choice, but also in the unfolding dynamics of the decision process itself (see also Devine et al., 2024, for effects of goal proximity on response caution in a cognitive task).

Taken together, these results suggest that progress toward a goal dynamically increases the subjective value of persistence, which is reflected both in choices and in the greater conflict that results from disengagement near the goal.

In addition to global progress effects, we observed an idiosyncratic dynamic with a pronounced impact of securing the first checkpoint on persistence decisions. The probability of disengagement dropped sharply after participants passed the first checkpoint, regardless of trial length. This abrupt shift suggests that initial progress plays an important role in establishing engagement, potentially by increasing perceived control or self-efficacy within the task (Shenhav et al., 2021a). However, because passing a checkpoint also reduces the remaining task and preserves future progress after failure, the present design does not isolate a purely motivational effect of early success from changes in task structure. Prior work indicates that early indicators of success can disproportionately influence motivation and subsequent effort allocation, as individuals infer their likelihood of success from initial performance cues (Carver and Scheier, 1990; Iso-Ahola and Dotson, 2014). Consistent with this interpretation, passing the first checkpoint may serve as a salient signal of control efficacy or reduced future cost, thereby increasing the expected value of continued effort investment (Shenhav et al., 2021a). Accordingly, once participants successfully passed the first checkpoint, they appeared more willing to continue investing effort, suggesting that clearing the first checkpoint may act as a gating signal for sustained persistence, perhaps acting as a motivational boost. An alternative explanation is that this effect reflects a form of loss aversion, whereby participants become unwilling to lose the benefit of having passed the first checkpoint. A further alternative is that this effect reflects sensorimotor learning concentrated early in each level. This account predicts that the effect should weaken with practice, but we found no evidence of such attenuation, even though performance improved over the experiment.

Beyond this initial transition, progress within trials exhibited different dynamics depending on task structure. To disentangle these effects, we compared equivalent checkpoint positions across trials of different lengths, allowing us to separate the influence of effort already invested from the effort remaining to reach the goal. When aligning checkpoints relative to the goal, participants were less likely to disengage in longer trials than in shorter ones. Because reaching a given distance from the goal required greater prior effort in longer trials, this pattern is consistent with a possible retrospective sensitivity to *sunk costs*, whereby previously invested effort increases commitment to the current course of action (Arkes and Blumer, 1985). However, this interpretation should be treated with caution, as the present design does not rule out alternative accounts, such as differences in relative progress framing or updated beliefs about successful completion. In addition, participants may have evaluated progress relative to total trial length, such that the same absolute distance represented greater relative proximity to the goal in longer trials (Devine and Otto, 2022).

In contrast, when checkpoints were aligned relative to the start of the trial, participants were more likely to disengage in longer than in shorter trials. Here, differences in remaining effort were held constant relative to progress, but the total effort required to complete the trial was greater in longer conditions. This pattern suggests that prospective evaluations of effort and delay reduced the subjective value of persistence, consistent with effort and delay discounting accounts (Westbrook et al., 2013; Prévost et al., 2010; Shenhav et al., 2017; Green and Myerson, 2004). Taken together, these findings indicate that persistence decisions are jointly shaped by retrospective and prospective evaluations of effort. Previously invested effort appears to increase commitment, whereas anticipated future effort reduces it.

More broadly, these results suggest that persistence decisions depend not only on absolute progress toward a goal, but also on how that progress is framed relative to reference points such as the start or the end of goal pursuit. By dynamically integrating signals about past investment and future demands, individuals appear to regulate persistence in a way that balances the costs and benefits of continued effort (Shenhav et al., 2013; Proietti et al., 2025).

Persistence decisions were also preceded by short-term drop in performance, but not in decision conflict. We observed a decline in performance across the final attempts preceding disengagement, indicating that decisions to disengage were typically preceded by a short sequence of poorer outcomes. This pattern suggests that recent performance provides a local signal influencing persistence decisions, consistent with the idea that individuals monitor ongoing success to evaluate whether continued effort is worthwhile (Carver and Scheier, 1990). This mechanism may also contribute to disengagement following performance decline, insofar as recent failures reduce inferred success probability and thus the expected value of persistence.

In contrast, decision dynamics did not show a corresponding change prior to disengagement. Mouse-tracking analyses revealed no systematic increase in trajectory curvature in the final attempts before disengagement, indicating that decision conflict did not progressively build up over these attempts (or that such a build-up is too subtle to be assessed in our setting). This pattern contrasts with the robust difference observed between persistence and disengagement decisions, with disengagement choices consistently associated with greater trajectory curvature, consistent with higher conflict at the moment of choice. This dissociation suggests that deterioration in performance influences the decision to disengage, without necessarily inducing an increase in the competition between response alternatives over preceding attempts. Rather, disengagement may reflect a sudden shift in decision-making strategy, in which the option to switch to easier settings emerges as a competitive alternative to the tendency to persist. However, this interpretation should be considered with caution, as persistence decisions were much more frequent, and greater familiarity with the persistence response may reduce apparent conflict for persistence choices and inflate observed differences between options. A similar imbalance can confound choice conflict with the value of switching in sequential decisions, a confound that joint modeling of choices and response times can help resolve (Fontanesi et al., 2022). Taken together, these findings indicate that persistence decisions are sensitive to recent performance outcomes, but that the decision to disengage may occur relatively abruptly rather than emerging from a gradual increase in internal conflict. This pattern is consistent with accounts in which disengagement is triggered when performance falls below a subjective threshold, rather than resulting from a continuous accumulation of decisional uncertainty.

Overall, these findings suggest that persistence decisions are governed by the dynamic integration of multiple signals operating at different timescales: global effort accumulation, progress (or lack of progress) toward the goal, and performance outcomes. These signals appear to converge on a common value-based decision process that determines whether to continue or disengage. This process may reflect a dynamic integration of effort costs and inferred success probability, whereby persistence is favored when the expected value of continued engagement outweighs the costs of exertion (Pessiglione et al., 2018).

Our results also add to a large literature showing that skilled and unskilled participants can adopt different decision strategies, plausibly as a result of differences in confidence regarding their ability to reach the goal, which may in turn influence cost–benefit computations, as well as other factors such as differences in farsightedness (Nuzzi et al., 2026; Moretti et al., 2025; Lancia et al., 2024; Coutrot et al., 2018; Krichmar and He, 2023; Lancia et al., 2023; Eluchans et al., 2025; van Opheusden et al., 2023). Importantly, the higher disengagement of unskilled participants should not be necessarily read as a motivational failure. Disengagement can be an adaptive response to the structure of the task rather than a lapse in motivation or self-control (McGuire and Kable, 2012, 2013): here, switching to the easier version of a level can be a reasonable strategy when the expected time and effort required to complete the harder version outweigh its additional reward, which is more often the case for less skilled participants. These findings underscore the importance of accounting for skill and other individual differences, particularly in studies involving naturalistic tasks, where substantial variability related to expertise, age, and other factors can be expected.

One limitation of this study is that we did not include direct physiological or experimental measures of fatigue. Although online data collection precluded the use of indices such as pupil size or EEG activity (van der Wel and van Steenbergen, 2018; Kurniawan et al., 2021; Cavanagh and Frank, 2014), the absence of an independent manipulation of fatigue also limits our ability to isolate its causal role. In addition, because participants could disengage at any time, fatigue accumulation may have been partially preempted by decisions to disengage, potentially restricting access to more extreme fatigue states. Future work should combine voluntary persistence paradigms with independent manipulations and physiological measures of fatigue.

Another limitation is that task incentives were held constant across conditions. The absence of a parametric manipulation of reward limits our ability to characterize how persistence decisions scale with expected value. Future work should vary reward magnitude and opportunity costs to more precisely examine how value signals interact with effort and progress.

Finally, individual differences were captured using a performance-based grouping approach. Although the main skill effects were replicated with a continuous measure of performance (see Appendix), both measures are derived from the task itself; future work could use independent measures of ability to more precisely characterize how ability interacts with effort and progress signals. Although the present task involved both motor and cognitive demands, the main source of effort likely lies in sustained attention and precise timing, suggesting a substantial cognitive effort component. Future research could extend this paradigm by systematically varying the balance between cognitive and motor demands, for example by increasing cognitive load (e.g., working memory requirements) or cognitive control demands, motor difficulty (Cools, 2016; Kool et al., 2010; Kool and Botvinick, 2018; Shenhav et al., 2017), to examine whether the observed persistence dynamics generalize across different forms and intensities of effort.

In sum, the present study provides a unified account of persistence in sequential effortful tasks, showing that decisions to persist or disengage are shaped by the interaction of effort costs, (lack of) progress, goal proximity, and early success. These findings highlight how goal-directed behavior is regulated over time, offering a framework for understanding its determinants. More broadly, this framework may provide a useful benchmark for characterizing alterations in persistence observed in clinical conditions. For instance, motivational deficits in depression have been linked to increased sensitivity to effort costs, rather than impairments in reward processing (Vinckier et al., 2022). The present paradigm may help disentangle whether such alterations reflect changes in effort valuation, reduced sensitivity to progress signals, or impaired integration of performance feedback. This would offer a practical approach to the computational phenotyping of persistence-related dysfunctions.

