## Supplementary Materials for "Determinants of persistence in sequential effort-based decision-making"

### Supplementary Materials: Determinants of persistence in sequential effort-based decision-making

### Appendix A Task and levels

#### A.1 Procedure for generating levels

Levels were procedurally generated by, for each level length $c \in \{\text{Short}, \text{Long}\}$ and level $l$, computing the total number of pipes ($\text{checkpoint\_target}_c \times \text{gap\_score}$), that is, $3 \times 4$ (Short) or $6 \times 4$ (Long). Each pipe was indexed, so that its horizontal position on the canvas corresponded directly to the index $i$.

The vertical position of each pipe opening was determined by drawing independent values from a uniform distribution:

$$y_i \sim \mathcal{U}\big(50,\ \text{canvas\_height} - \text{gap} - 100\big),$$

where canvas_height = 770 px. Difficulty was modulated by adapting the size of the vertical gap between pipes: in D1 levels the gap was relatively large (gap = 310 px), whereas in D2 levels the gap was reduced (gap = 240 px), and by adapting the horizontal spacing between consecutive pipes (D1: 310 px; D2: 290 px). A smaller gap increases the precision required from the player, thereby raising task difficulty. The easier version of each level (D0) used larger vertical and horizontal gaps than both D1 and D2 (vertical gap: 560 px; horizontal spacing: 410 px).

The top pipe extended from ($x = i$, $y = y_i + \text{gap}$) up to canvas_height, while the bottom pipe extended from the base of the canvas ($y = 0$) up to $y = y_i$. Checkpoints were deterministically assigned every fourth pipe.

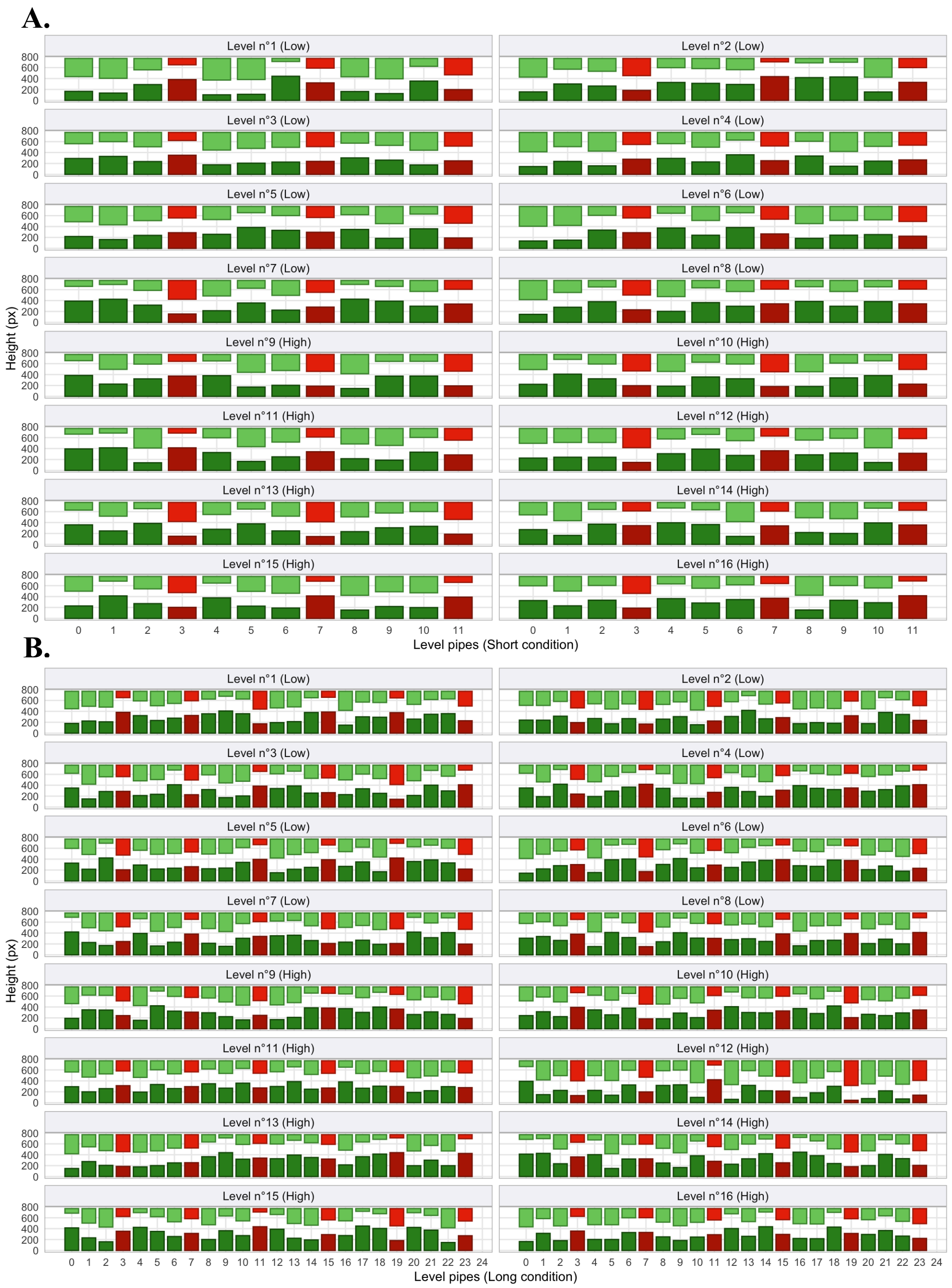


**Fig. A1** Procedurally generated levels by length and difficulty. Each panel shows one level; bars show the bottom (dark green) and top (light green) pipes at each pipe position (x-axis), and the gap between them is the opening the bird must fly through (y-axis: height in pixels). Red pipes indicate checkpoints. “Low” and “High” denote D1 (easier) and D2 (harder) levels, respectively. (A) Short (3 CP) levels. (B) Long (6 CP) levels.

#### A.2 Number of attempts per level

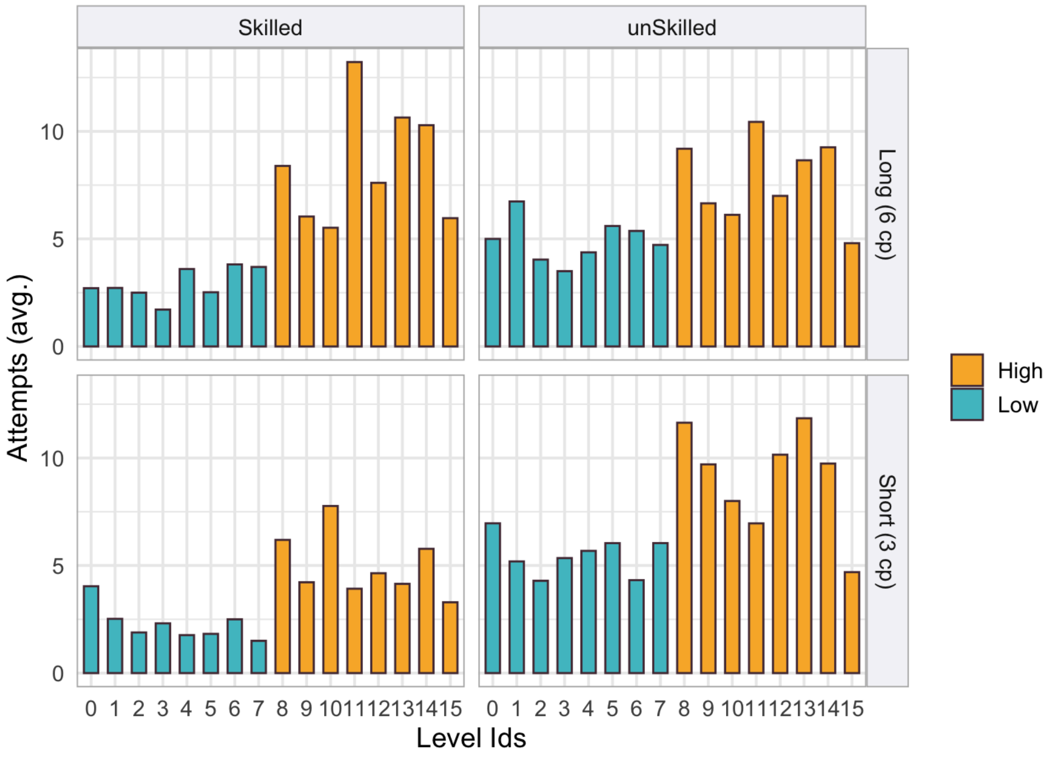


**Fig. A2** Mean number of attempts per level, by level length (rows: Long, 6 CP; Short, 3 CP), skill group and difficulty ("Low" = D1, easier; "High" = D2, harder). Level IDs (0–15) index the 16 levels within each level length; IDs 0–7 are D1 and 8–15 are D2 levels.

### Appendix B  Mouse-tracking preprocessing

Mouse cursor trajectories were recorded continuously during the choice phase at a nominal sampling rate of 10 ms. Trajectory preprocessing followed standard procedures in mouse-tracking research [1, 2] and was implemented prior to feature extraction.

##### *Trajectory alignment*

To standardize trajectories across participants, all trajectories were remapped to a common coordinate frame. For each participant, we first estimated the median valid starting position across trials and the median temporal gap between stimulus onset and movement initiation. All trajectories were then translated such that their first valid point corresponded to the coordinate origin $(0, 0)$. Because the two response options were located on opposite sides of the screen, trajectories corresponding to "Make it easier" responses were mirrored along the x-axis so that all trajectories reflected movement toward the same side. This alignment allowed aggregation and comparison of trajectories independently of response side.

##### *Coordinate system*

The game interface was identical across participants and trials. Consequently, trajectory coordinates were analyzed in raw pixel space and were not rescaled.

##### *Trajectory exclusion*

Trajectories were excluded if:

- the final cursor position was abnormally low relative to the response region,
- movement initiation latency exceeded 1000 ms,
- temporal anomalies indicated irregular or discontinuous cursor recording.

Across all recorded attempts ($N$ = 9,434), 522 trajectories (5.5%) met one or more of these criteria. A further 666 trajectories whose sampling interval deviated from the nominal 10 ms were removed. The final dataset comprised 8,246 trajectories (87.4% of attempts).

##### *Time normalization and feature extraction*

All valid trajectories were time-normalized to 101 equally spaced time steps using linear interpolation, which preserves trajectory shape while aligning movements of different durations to a common temporal scale. Deviation profiles were smoothed using a centered moving-average filter with a 7-point window. For each trajectory, we extracted the maximal absolute deviation ($MAD$), that is, the maximum perpendicular deviation from the ideal straight line between the starting point and the selected response option. $MAD$ indexes the degree of attraction toward the unchosen alternative and is commonly interpreted as an indicator of decision conflict.

### Appendix C Descriptive statistics

#### C.1 Scores and disengagement rates

Final scores ranged from 32 to 160 points ($M = 103$, $SD = 48.2$; skilled: $M = 147$, $SD = 13.4$; unskilled: $M = 59.6$, $SD = 24.2$).

Participants made 9,434 decisions to persist or disengage, of which 713 (7.6%) were decisions to disengage. Table C1 reports disengagement rates at two levels: per decision (the proportion of persist/disengage decisions that were disengagements) and per level (the proportion of levels completed in their easier version). The two measures can diverge: harder levels produce more failed attempts, and thus many more persist decisions, so the per-decision rate can be lower in harder levels even when more of these levels end in disengagement. The model estimates (main text, Table 1) are per decision and adjust for the other predictors.

**Table C1** Disengagement rates by difficulty and level length, and by skill group.

| | | Per decision | | Per level (% of levels) | | |
|---|---|---|---|---|---|---|
| Difficulty | Level length | $n$ | % disengage | $n_{\text{participants}}$ | $M$ | $SD$ |
| D1 (easier) | Long (6 CP) | 1,586 | 10.0 | 56 | 36.2 | 43.1 |
| D1 (easier) | Short (3 CP) | 1,445 | 8.2 | 56 | 27.0 | 36.7 |
| D2 (harder) | Long (6 CP) | 3,447 | 6.7 | 56 | 53.5 | 40.7 |
| D2 (harder) | Short (3 CP) | 2,956 | 6.9 | 56 | 46.5 | 42.9 |
| Skilled | | | | 28 | 20.3 | 25.7 |
| Unskilled | | | | 28 | 60.7 | 36.3 |

*Note.* Per-decision rates are pooled across participants. Per-level rates were computed for each participant and then averaged; individual rates ranged from 0% to 100%.

#### C.2 Grit

Grit was measured with the Short Grit Scale [3]. We tested whether grit scores were associated with each participant's overall disengagement rate and with skill (mean number of obstacles passed per attempt, standardized). Neither association was significant (Table C2).

**Table C2** Correlations between grit and task behavior ($N = 56$).

| | Spearman | | | Pearson | | |
|---|---|---|---|---|---|---|
| | $\rho$ | 95% CI | $p$ | $r$ | $p$ | $p_{\text{holm}}$ |
| Disengagement rate | .06 | [−.24, .36] | .699 | .19 | .233 | 1 |
| Skill | −.02 | [−.32, .28] | .906 | −.01 | .933 | 1 |

### Appendix D Skill groups

#### D.1 Definition and continuous measure

For each participant, skill was computed as the mean number of obstacles passed per attempt in D1 and D2 levels. Participants were split at the median (2.34 obstacles per attempt) into a skilled and an unskilled group ($n = 28$ each). For the continuous analyses, the same quantity was standardized across participants. The two codings were closely related (point-biserial $r = .76$, $t(54) = 8.71$, $p < .001$). Because both measures are derived from task performance, skill is a performance-based descriptor rather than an independent individual-difference measure.

#### D.2 Checkpoint progression as a function of skill group

To describe how the two groups progressed through the levels, we analyzed within-attempt checkpoint advancement using a linear mixed-effects model. For each attempt, progress was defined as the difference between the ending and the starting checkpoint. Progress was modeled as a function of *skill group*, *starting checkpoint*, *level length* and their interactions, and *difficulty*, with a random intercept for participant.

As expected given how the groups were defined, unskilled participants progressed less per attempt than skilled participants ($\beta = -0.823$, $SE = 0.080$, $t = -10.31$, $p < .001$). Progress decreased with the starting checkpoint ($\beta = -0.120$, $SE = 0.008$, $t = -15.29$, $p < .001$), reflecting the shorter remaining distance when attempts began closer to the goal. The interaction between skill group and starting checkpoint was significant ($\beta = 0.108$, $SE = 0.011$, $t = 9.76$, $p < .001$), indicating that differences between groups were largest at earlier checkpoints. Progress was greater in easier levels ($\beta = 0.528$, $SE = 0.015$, $t = 34.34$, $p < .001$) and slightly lower in short than in long levels ($\beta = -0.226$, $SE = 0.032$, $t = -7.06$, $p < .001$). The three-way interaction between skill group, starting checkpoint and level length was not significant ($p = .719$).

#### D.3 Disengagement model with continuous skill

To test whether the skill-related results depend on the median split, we fitted the disengagement model with skill as a continuous predictor. Its fixed-effects structure was selected with the same forward LRT procedure as the main model; the interaction between distance from goal and skill did not improve fit and was not retained. The model used the same random-effects structure as the main model (Appendix E.2).

Table D3 compares the fit of the four specifications, and Table D4 reports the fixed effects of the continuous-skill model with random slopes. All skill-related effects of the main model were replicated. Coefficients for skill have the opposite sign to those of the binary coding, because the continuous score increases with skill whereas the binary predictor codes the unskilled group.

**Table D3** Model fit across random-effects structures and skill codings.

| Model | AIC | BIC | ΔAIC |
|---|---|---|---|
| Binary skill, random intercept only (submitted model) | 3377.9 | 3492.3 | — |
| Binary skill, revised random effects (main model) | 3092.7 | 3257.2 | −285.1 |
| Continuous skill, random intercept only | 3379.5 | 3486.8 | +1.7 |
| Continuous skill, revised random effects | 3101.3 | 3258.7 | −276.6 |

*Note.* ΔAIC is relative to the submitted model. All models were fitted to 9,434 decisions from 56 participants.

**Table D4** Fixed-effect estimates of the disengagement model with continuous skill and revised random effects (logit link).

| Predictor | Estimate | *SE* | *z* | *p* | 95% CI |
|---|---|---|---|---|---|
| Intercept | −3.964 | 0.331 | −11.97 | $< .001$ | [−4.612, −3.315] |
| Distance from goal | 1.110 | 0.149 | 7.43 | $< .001$ | [0.817, 1.403] |
| Skill (continuous) | −1.770 | 0.360 | −4.92 | $< .001$ | [−2.474, −1.065] |
| Level difficulty D1 (vs. D2[a]) | −0.317 | 0.119 | −2.66 | $< .01$ | [−0.550, −0.083] |
| Attempts (poly$^1$) | 47.275 | 13.247 | 3.57 | $< .001$ | [21.311, 73.239] |
| Attempts (poly$^2$) | 17.686 | 5.878 | 3.01 | $< .01$ | [6.165, 29.207] |
| Obstacles passed (poly$^1$) | −17.392 | 8.159 | −2.13 | $< .05$ | [−33.384, −1.400] |
| Obstacles passed (poly$^2$) | −6.488 | 6.532 | −0.99 | .321 | [−19.292, 6.315] |
| Trial index (poly$^1$) | −7.416 | 13.842 | −0.54 | .592 | [−34.546, 19.715] |
| Trial index (poly$^2$) | −5.169 | 6.194 | −0.83 | .404 | [−17.309, 6.970] |
| Skill × Attempts (poly$^1$) | 57.657 | 13.374 | 4.31 | $< .001$ | [31.446, 83.869] |
| Skill × Attempts (poly$^2$) | −30.676 | 6.675 | −4.60 | $< .001$ | [−43.758, −17.594] |
| Skill × Trial index (poly$^1$) | −22.886 | 14.648 | −1.56 | .118 | [−51.595, 5.823] |
| Skill × Trial index (poly$^2$) | 28.520 | 6.892 | 4.14 | $< .001$ | [15.012, 42.028] |

*Note.* Model: Pr(Disengagement) ∼ distance to goal + skill + difficulty + poly(*attempt index*, 2) + poly(*obstacles passed*, 2) + poly(*trial index*, 2) + skill × poly(*attempt index*, 2) + skill × poly(*trial index*, 2) + (1 + distance to goal + *attempt index* | participant) + (0 + *trial index* || participant) + (0 + *obstacles passed* || participant). Skill is the standardized mean number of obstacles passed per attempt. Estimates and 95% Wald CIs are on the log-odds scale. [a] Reference category.

### Appendix E Disengagement model

#### E.1 Fixed-effect selection

Disengagement was modeled with binomial generalized linear mixed-effects models (GLMMs) with a logit link. The fixed-effects structure was selected first, with a random intercept for participant. The base model included *distance from goal*, *skill group*, *difficulty*, and quadratic polynomial terms for *attempt index*, *obstacles passed* and *trial index*. Interaction terms were then added and retained if they improved fit in likelihood-ratio tests (LRTs).

Table E5 summarizes the candidate models and Table E6 their fit. Adding the interaction between *skill group* and *attempt index* (Model 003) substantially improved fit relative to the base model. The interaction between *distance from goal* and *skill group* did not improve fit when added to the base model (Model 002), but did when added to Model 003 (Model 007). Adding the interaction between *skill group* and *trial index* further improved fit (Model 008), which had the lowest AIC and BIC and was retained. The random-effects structure of this model was then selected as described in Appendix E.2.

**Table E5** Fixed effects of the candidate models.

| Model | Fixed effects |
|---|---|
| 001 | Distance from goal + Skill group + Difficulty + poly(Attempt index, 2) + poly(Obstacles passed, 2) + poly(Trial index, 2) |
| 002 | 001 + Distance from goal × Skill group |
| 006 | 001 + Distance from goal × Difficulty |
| 003 | 001 + Skill group × poly(Attempt index, 2) |
| 004 | 001 + Skill group × poly(Obstacles passed, 2) |
| 005 | 001 + Skill group × poly(Trial index, 2) |
| 007 | 003 + Distance from goal × Skill group |
| 008 | 007 + Skill group × poly(Trial index, 2) |

#### E.2 Random-effects selection

Following recommendations to model by-participant variability in within-subject effects [4] while retaining only the random-effects structure supported by the data, we started from the selected fixed-effects model with a random intercept for participant (Model 008) and tested a by-participant random slope

**Table E6** Fit of the candidate models (random intercept for participant).

| Model | npar | AIC | BIC | logLik | Deviance | Compared with | $\chi^2$ (df) | $p$ |
|---|---|---|---|---|---|---|---|---|
| 001 | 11 | 3470.3 | 3549.0 | −1724.2 | 3448.3 | — | — | — |
| 002 | 12 | 3472.3 | 3558.2 | −1724.2 | 3448.3 | 001 | 0.01 (1) | .926 |
| 006 | 12 | 3470.8 | 3556.6 | −1723.4 | 3446.8 | 001 | 1.5 (1) | .221 |
| 003 | 13 | 3416.5 | 3509.5 | −1695.2 | 3390.5 | 001 | 57.8 (2) | < .001 |
| 004 | 13 | 3468.6 | 3561.6 | −1721.3 | 3442.6 | 001 | 5.7 (2) | .058 |
| 005 | 13 | 3429.0 | 3522.0 | −1701.5 | 3403.0 | 001 | 45.3 (2) | < .001 |
| 007 | 14 | 3414.4 | 3514.5 | −1693.2 | 3386.4 | 003 | 4.1 (1) | < .05 |
| 008 | 16 | **3377.9** | **3492.3** | −1672.9 | 3345.9 | 007 | 40.5 (2) | < .001 |

for each within-subject predictor: distance from goal, attempt index, obstacles passed, trial index and difficulty. Each slope was first added on its own; slopes were then combined, allowed to correlate with the intercept where this improved fit, and each remaining candidate was re-tested against the combined structure (Table E7). Slopes were retained if they improved fit in LRTs without producing a singular fit. The final structure comprised a random intercept correlated with random slopes for distance from goal and attempt index, and uncorrelated random slopes for trial index and obstacles passed. The slope for difficulty did not improve fit and was not retained. Overall, the final structure improved fit substantially relative to the intercept-only model ($\chi^2(7) = 299.14$, $p < .001$; $\Delta\text{AIC} = -285.1$).

**Table E7** Selection of the random-effects structure of the disengagement model.

| Model | Random effects | npar | AIC | BIC | logLik | Compared with | $\chi^2$ (df) | $p$ |
|---|---|---|---|---|---|---|---|---|
| r00 | (1 \| participant) | 16 | 3377.9 | 3492.3 | −1672.9 | — | — | — |
| *Single slopes* | | | | | | | | |
| r01 | r00 + distance | 17 | 3256.6 | 3378.2 | −1611.3 | r00 | 123.25 (1) | < .001 |
| r02 | r00 + attempt | 17 | 3300.8 | 3422.4 | −1633.4 | r00 | 79.04 (1) | < .001 |
| r04 | r00 + obstacles | 17 | 3371.3 | 3492.8 | −1668.6 | r00 | 8.60 (1) | < .01 |
| r05 | r00 + trial | 17 | 3188.9 | 3310.5 | −1577.4 | r00 | 190.96 (1) | < .001 |
| r03 | r00 + difficulty | 19 | 3376.1 | 3512.0 | −1669.0 | r00 | 7.77 (3) | .051 |
| *Combined structure* | | | | | | | | |
| r06 | r01 + attempt (uncorrelated) | 18 | 3188.0 | 3316.8 | −1576.0 | r01 | 70.56 (1) | < .001 |
| r07 | r06, correlated with intercept | 21 | 3165.9 | 3316.1 | −1561.9 | r06 | 28.16 (3) | < .001 |
| r12 | r07 + trial | 22 | 3098.3 | 3255.6 | −1527.1 | r07 | 69.62 (1) | < .001 |
| **r13** | **r12 + obstacles** | 23 | **3092.7** | 3257.2 | −1523.4 | r12 | 7.56 (1) | < .01 |
| r14 | r12 + difficulty | 25 | 3102.6 | 3281.4 | −1526.3 | r12 | 1.64 (3) | .650 |

*Note.* All models share the fixed effects of Model 008. "Distance", "attempt", "obstacles", "trial" and "difficulty" denote by-participant random slopes for distance from goal, attempt index, obstacles passed, trial index and difficulty. None of the models was singular. The final model (r13) is the model reported in the main text (Table 1).

#### E.3 Final model: random effects and diagnostics

The final model was fitted to 9,434 decisions from 56 participants (AIC = 3092.7, BIC = 3257.2, $\log L = -1523.4$). Random-effect standard deviations were 2.69 for the intercept, 0.68 for distance from goal, 0.83 for attempt index, 0.97 for trial index and 0.27 for obstacles passed.

Multicollinearity was assessed using generalized variance inflation factors (GVIFs). All adjusted GVIF values were below 2 (Table E8), indicating no problematic multicollinearity among predictors.

#### E.4 Failure likelihood as a predictor of disengagement

To test whether the effects of skill and difficulty on disengagement could be summarized by the likelihood of failing, we proceeded in two steps. First, we modeled the probability of failing at each obstacle as a function of skill group, difficulty and their interaction, using a logistic model fitted to all obstacles encountered during D1 and D2 attempts (with a random intercept for participant). Failure was more likely for unskilled participants ($\beta = 1.08$, $p < .001$) and less likely in easier levels ($\beta = -1.07$, $p < .001$),

**Table E8** Generalized variance inflation factors for the predictors of the final model.

| Predictor | df | $\mathrm{GVIF}^{1/(2\,\mathrm{df})}$ | |
|---|---|---|---|
| Distance from goal | 3.87 | 1 | 1.97 |
| Skill group | 1.79 | 1 | 1.34 |
| Level difficulty (D1 vs. D2) | 1.03 | 1 | 1.02 |
| Attempts (poly, degree 2) | 7.32 | 2 | 1.64 |
| Obstacles passed (poly, degree 2) | 1.03 | 2 | 1.01 |
| Trial index (poly, degree 2) | 4.84 | 2 | 1.48 |
| Skill × Attempts (poly, degree 2) | 9.77 | 2 | 1.77 |
| Distance × Skill group | 3.85 | 1 | 1.96 |
| Skill × Trial index (poly, degree 2) | 4.85 | 2 | 1.48 |

*Note.* GVIFs were computed with `car::vif`.

with a significant interaction ($\beta = 0.32$, $p < .001$): the reduction in failure risk from harder to easier levels was smaller for unskilled participants. Because skill groups were defined by performance, the skill effect in this model is expected by construction. Second, we used this model to compute, for each attempt of the main dataset, a predicted failure likelihood based only on skill group and difficulty, and entered it in the disengagement model in place of skill and difficulty.

A model with failure likelihood alone did not fit worse than a model including skill, difficulty and their interaction (Table E9, rows 1–2). When the interactions of skill with attempt index and trial index were included, the specification with separate predictors fit somewhat better on AIC and similarly on BIC (rows 3–4); these two models are not nested, so this comparison is descriptive only. Because failure likelihood takes only four values (two skill groups × two difficulty levels), these results show that a single failure-likelihood predictor is an adequate summary of the main effects of skill and difficulty, rather than strong evidence for a single mechanism.

**Table E9** Comparison of models with skill and difficulty or with failure likelihood as predictors of disengagement.

| | Model | AIC | BIC | Test |
|---|---|---|---|---|
| 1 | Skill × difficulty (main effects) | 3154.8 | 3290.7 | — |
| 2 | Failure likelihood (main effect) | 3151.9 | 3273.5 | $\chi^2(2) = 1.11$, $p = .575$ vs. (1) |
| 3 | Skill + difficulty (with interactions) | 3092.7 | 3257.2 | — |
| 4 | Failure likelihood (with interactions) | 3101.8 | 3259.1 | not nested with (3) |

*Note.* All models include the other fixed effects and the random-effects structure of the main model. Rows 3–4 additionally include the interactions of skill (or failure likelihood) with attempt index and trial index, as in the main model.

#### E.5 Auxiliary analyses

Two auxiliary analyses assessed the robustness of the main model. Both were fitted with the fixed and random effects of the final model.

##### *Between-level variance*

To assess potential stimulus-sampling effects, we added a random intercept for level (32 levels) to the final model. The between-level variance was small relative to the between-participant variance (SD = 0.54 vs. 2.66; Table E10).

##### *Block order*

To test whether the order of the two blocks (short-first vs. long-first) influenced disengagement, we added block order as a fixed effect to the final model. Block order did not significantly predict disengagement (short-first vs. long-first: $\beta = 0.17$, $SE = 0.57$, $z = 0.29$, $p = .770$) and did not improve fit ($\chi^2(1) = 0.08$, $p = .771$); all other effects kept their direction and significance, with nearly identical estimates.

**Table E10** Variance components of the final model with an additional random intercept for level.

| Grouping factor | Effect | Variance | SD |
|---|---|---|---|
| Participant | Intercept | 7.064 | 2.658 |
| | Distance from goal | 0.534 | 0.731 |
| | Attempt index | 0.676 | 0.822 |
| | Trial index | 0.806 | 0.898 |
| | Obstacles passed | 0.080 | 0.283 |
| Level | Intercept | 0.287 | 0.535 |

*Note.* Fixed effects kept their direction relative to the final model (main text, Table 1); level difficulty was no longer significant ($\beta = -0.33$, $SE = 0.23$, $p = .139$).

### Appendix F Checkpoint-specific model

#### F.1 Model specification

Because *level length* and *distance from goal* are structurally collinear, the main model treated *distance from goal* as a continuous predictor. To test for checkpoint-specific deviations from this global trend, we fitted a GLMM in which distance was coded as a categorical factor indexing checkpoint position within each level length (nine levels), with its interaction with skill group as fixed effects. A full by-participant random slope for this nine-level factor was not feasible with the present data. We therefore tested random slopes for the two continuous dimensions on which the planned contrasts are built, distance from goal and level length (Table F11). Both improved fit, and the model with both slopes was retained; it was preferred on AIC and the LRT, and was close to the distance-only model on BIC. Planned contrasts were computed with *emmeans* and Holm-corrected within each family of contrasts.

**Table F11** Selection of the random-effects structure of the checkpoint-specific model.

| Random effects | npar | AIC | BIC | logLik | Compared with | $\chi^2$ (df) | $p$ |
|---|---|---|---|---|---|---|---|
| (a) (1 \| participant) | 19 | 3480.3 | 3616.2 | −1721.2 | — | — | — |
| (b) (1 + distance \| participant) | 21 | 3287.5 | 3437.7 | −1622.7 | (a) | 196.87 (2) | < .001 |
| **(c) (1 + distance + level length \| participant)** | 24 | **3268.9** | 3440.6 | −1610.5 | (b) | 24.57 (3) | < .001 |

*Note.* Fixed effects: Pr(Disengagement) ∼ distance by condition + distance by condition × skill. None of the models was singular. Model (c) is the model reported in the main text (Table 2).

#### F.2 Planned contrasts

Table F12 reports the complete set of planned contrasts; the significant ones are shown in the main text (Table 2).

#### F.3 Learning and the first-checkpoint effect

If the drop in disengagement after the first checkpoint reflected sensorimotor learning, it should attenuate as participants become more practiced. We first tested whether performance improved over the experiment, using a Poisson GLMM predicting the number of obstacles passed per attempt from a quadratic polynomial of trial index with a random intercept for participant. Performance improved over trials (linear trend: $\beta = 11.78$, $SE = 0.72$, $p < .001$), with a decelerating trajectory (quadratic trend: $\beta = -5.03$, $SE = 0.67$, $p < .001$).

We then fitted, separately for long and short levels, a GLMM restricted to the two checkpoints that define the first-checkpoint transition (long: 6CP vs. 5CP; short: 3CP vs. 2CP), predicting disengagement from checkpoint, block position (first vs. second block), their interaction and difficulty, with a random intercept for participant. Because level length was blocked, block position compares participants who

**Table F12** Planned contrasts comparing disengagement probabilities between short and long levels at matched distances from the goal and from the start, and between consecutive checkpoints within each level length.

| Contrast | Odds ratio | *SE* | $z$ | $p_{\text{holm}}$ |
|---|---|---|---|---|
| *All participants — distance from goal* | | | | |
| Short vs Long (3CP) | 13971.41 | 64167.70 | 2.08 | .075 |
| Short vs Long (2CP) | 1.81 | 1.16 | 0.93 | .350 |
| Short vs Long (1CP) | 2719.12 | 4265.21 | 5.04 | $< .001$ |
| *All participants — distance from start* | | | | |
| Short vs Long (Start, CP0) | 0.37 | 0.132 | −2.79 | $< .05$ |
| Short vs Long (CP1) | 0.69 | 0.303 | −0.85 | .396 |
| Short vs Long (CP2) | 0.29 | 0.142 | −2.53 | $< .05$ |
| *Within level length — consecutive checkpoints* | | | | |
| Long 6CP vs 5CP | 4.73 | 1.179 | 6.23 | $< .001$ |
| Long 5CP vs 4CP | 0.69 | 0.229 | −1.12 | .261 |
| Long 4CP vs 3CP | 11713.35 | 53785.41 | 2.04 | .124 |
| Long 3CP vs 2CP | 0.00033 | 0.00151 | −1.74 | .164 |
| Long 2CP vs 1CP | 2421.60 | 3937.43 | 4.79 | $< .001$ |
| Short 3CP vs 2CP | 2.52 | 0.600 | 3.87 | $< .001$ |
| Short 2CP vs 1CP | 1.62 | 0.568 | 1.37 | .172 |
| *By skill group — distance from goal* | | | | |
| Short vs Long (3CP) — Skilled | 1.35e+07 | 1.22e+08 | 1.81 | .281 |
| Short vs Long (2CP) — Skilled | 0.65 | 0.435 | −0.65 | 1 |
| Short vs Long (1CP) — Skilled | 5.52e+06 | 1.66e+07 | 5.17 | $< .001$ |
| Short vs Long (3CP) — Unskilled | 14.50 | 15.30 | 2.53 | .057 |
| Short vs Long (2CP) — Unskilled | 5.08 | 5.50 | 1.50 | .401 |
| Short vs Long (1CP) — Unskilled | 1.34 | 1.17 | 0.33 | 1 |
| *By skill group — distance from start* | | | | |
| Short vs Long (Start, CP0) — Skilled | 0.34 | 0.191 | −1.92 | .272 |
| Short vs Long (CP1) — Skilled | 0.45 | 0.291 | −1.23 | .435 |
| Short vs Long (CP2) — Skilled | 0.30 | 0.206 | −1.75 | .272 |
| Short vs Long (Start, CP0) — Unskilled | 0.39 | 0.157 | −2.34 | .117 |
| Short vs Long (CP1) — Unskilled | 1.05 | 0.593 | 0.09 | .930 |
| Short vs Long (CP2) — Unskilled | 0.29 | 0.189 | −1.89 | .272 |

*Note.* Pairwise contrasts from the model in Table F11 (c); $z$ and $p$ values are computed on the log-odds scale and Holm-corrected within each family of contrasts. Extreme odds ratios reflect sparse disengagements in some cells and should be interpreted with caution.

encountered a given level length first with those who encountered it second. The checkpoint × block position interaction was not significant in either level length, and the drop in disengagement was not smaller in the second block (Table F13).

**Table F13** Predicted probability of disengagement at the first-checkpoint transition, by block position.

| Level length | Checkpoint | Block position | $P$(disengage) | 95% CI |
|---|---|---|---|---|
| Long | 6CP | First | .144 | [.059, .310] |
| Long | 6CP | Second | .091 | [.033, .224] |
| Long | 5CP | First | .029 | [.010, .081] |
| Long | 5CP | Second | .013 | [.004, .046] |
| Short | 3CP | First | .056 | [.022, .135] |
| Short | 3CP | Second | .058 | [.023, .138] |
| Short | 2CP | First | .018 | [.006, .052] |
| Short | 2CP | Second | .018 | [.006, .050] |

*Note.* Checkpoint × block position interaction: long levels, $\chi^2(1) = 0.25$, $p = .620$; short levels, $\chi^2(1) = 0.07$, $p = .798$ (both $p_{\text{holm}} = 1$).

### Appendix G Performance before disengagement

To characterize how performance evolved before disengagement, we analyzed the three attempts preceding the attempt that ended in a decision to disengage (attempt relative to disengagement: −3, −2, −1). The attempt that ended in disengagement was not included. The analysis comprised 754 attempts from 44 participants. Performance (number of obstacles passed) was modeled using a GLMM with a Poisson distribution and log link, including attempt relative to disengagement (entered as an ordered factor, with linear and quadratic trends), skill group, level length and all their interactions, with a random intercept for participant.

The number of obstacles passed decreased across the three attempts (linear trend: $\beta = -0.31$, $SE = 0.11$, $z = -2.88$, $p < .01$). Unskilled participants passed fewer obstacles than skilled participants ($\beta = -0.72$, $SE = 0.17$, $z = -4.16$, $p < .001$), and performance was lower in short than in long levels ($\beta = -0.33$, $SE = 0.11$, $z = -3.13$, $p < .01$). No interaction involving attempt relative to disengagement was significant (all $p$s $> .17$), indicating that the decline was comparable across skill groups and level lengths (Table G14).

**Table G14** Poisson GLMM predicting the number of obstacles passed in the three attempts preceding disengagement.

| Predictor | Estimate | $SE$ | $z$ | $p$ |
|---|---|---|---|---|
| Intercept | 0.582 | 0.127 | 4.60 | < .001 |
| Attempt (linear) | −0.309 | 0.107 | −2.88 | < .01 |
| Attempt (quadratic) | −0.160 | 0.099 | −1.62 | .106 |
| Unskilled (vs. Skilled[a]) | −0.720 | 0.173 | −4.16 | < .001 |
| Short (vs. Long[a]) | −0.328 | 0.105 | −3.13 | < .01 |
| Attempt (linear) × Unskilled | 0.224 | 0.166 | 1.35 | .177 |
| Attempt (quadratic) × Unskilled | 0.096 | 0.157 | 0.61 | .543 |
| Attempt (linear) × Short | 0.183 | 0.178 | 1.03 | .304 |
| Attempt (quadratic) × Short | −0.004 | 0.162 | −0.02 | .981 |
| Unskilled × Short | 0.290 | 0.149 | 1.94 | .052 |
| Attempt (linear) × Unskilled × Short | −0.276 | 0.247 | −1.12 | .263 |
| Attempt (quadratic) × Unskilled × Short | −0.080 | 0.230 | −0.35 | .729 |

*Note.* Model: obstacles passed ∼ attempt relative to disengagement × skill × level length + (1 | participant), Poisson distribution, log link. "Attempt" denotes attempt relative to disengagement, coded as an ordered factor with orthogonal polynomial contrasts over −3, −2, −1; negative linear coefficients indicate lower performance closer to disengagement. [a] Reference category.

### Appendix H Decision dynamics

#### H.1 Model specification and random-effects selection

Log-transformed *MAD* was modeled with a linear mixed-effects model including *decision* (persist vs. disengage), *distance from goal* and their interaction, with *skill group*, *difficulty*, *attempt index*, *trial index* and *obstacles passed* as control predictors. Nonlinear (quadratic) terms for attempt index, trial index and obstacles passed did not improve fit and were not retained.

The random-effects structure was selected with the same procedure as for the disengagement model (Table H15). The final structure improved fit substantially relative to the intercept-only model ($\Delta$AIC $= -199$).

**Table H15** Selection of the random-effects structure of the mouse-tracking model.

| Model | Random effects | npar | AIC | BIC | Compared with | $\chi^2$ (df) | *p* |
|---|---|---|---|---|---|---|---|
| m0 | (1 \| participant) | 11 | 17937 | 18014 | — | — | — |
| *Single slopes* | | | | | | | |
| m1 | (1 + decision \| participant) | 13 | 17809 | 17900 | m0 | 131.78 (2) | $< .001$ |
| m2 | m0 + trial | 12 | 17876 | 17960 | m0 | 62.60 (1) | $< .001$ |
| m3 | m0 + distance | 12 | 17918 | 18003 | m0 | 20.39 (1) | $< .001$ |
| m4 | m0 + attempt | 12 | 17926 | 18010 | m0 | 13.05 (1) | $< .001$ |
| m5 | m0 + obstacles | 12 | 17927 | 18011 | m0 | 12.12 (1) | $< .001$ |
| m6 | (1 + difficulty \| participant) | 13 | 17938 | 18029 | m0 | 3.33 (2) | .190 |
| *Combined structure* | | | | | | | |
| m7 | m1 + trial | 14 | 17748 | 17846 | m1 | 63.41 (1) | $< .001$ |
| m8 | m7 + obstacles | 15 | 17740 | 17845 | m7 | 9.59 (1) | $< .01$ |
| **m9** | **m8 + distance** | 16 | **17738** | 17850 | m8 | 4.42 (1) | $< .05$ |
| m10 | m9 + attempt | 17 | 17737 | 17856 | m9 | 2.83 (1) | .092 |

*Note.* All models share the same fixed effects and were compared on their REML fits. The final model (m9) is the model reported in the main text (Table 3).

**Table H16** Simple slopes of distance from goal on log-transformed *MAD*, by decision.

| Decision | Slope | *SE* | *z* | *p* |
|---|---|---|---|---|
| Disengage | −0.085 | 0.035 | −2.43 | $< .05$ |
| Persist | 0.007 | 0.013 | 0.59 | .554 |

*Note.* Slopes computed with *emtrends*, averaged over skill group and difficulty (asymptotic degrees of freedom). Negative slopes indicate greater curvature closer to the goal.

#### H.2 Effects of skill

Skill group was included as a covariate in the model. To test whether skill moderated the two focal effects, we added the interactions of skill group with decision type and with distance from goal to the final model, separately and together, and compared models with LRTs (maximum-likelihood fits). Skill did not predict trajectory curvature and did not moderate either effect (Table H17). We did not test the three-way interaction between skill, decision and checkpoint, because skilled participants rarely disengaged and several of the corresponding cells contained few trajectories.

#### H.3 Decision dynamics before disengagement

To test whether decision conflict increased before disengagement, we analyzed log-transformed *MAD* across the three attempts preceding the attempt that ended in a decision to disengage, coded as in Appendix G. The model included attempt relative to disengagement (ordered factor, linear and quadratic

**Table H17** Effects of skill group on trajectory curvature.

| Effect | Estimate / $\chi^2$ | df | $p$ |
|---|---|---|---|
| Skill (main effect) | $\beta = -0.079$ ($SE = 0.083$) | 46.4 | .344 |
| Skill × Decision | $\chi^2 = 1.33$ | 1 | .249 |
| Skill × Distance from goal | $\chi^2 = 0.81$ | 1 | .369 |
| Both interactions | $\chi^2 = 2.02$ | 2 | .364 |

trends), skill group and their interaction, with a random intercept for participant. None of the predictors significantly affected trajectory curvature (Table H18), indicating no detectable change in decision conflict over these attempts.

**Table H18** Linear mixed-effects model predicting log-transformed $MAD$ across the three attempts preceding disengagement.

| Predictor | Estimate | $SE$ | $t$ | $p$ |
|---|---|---|---|---|
| Intercept | 3.744 | 0.102 | 36.85 | $< .001$ |
| Attempt (linear) | 0.107 | 0.098 | 1.09 | .276 |
| Attempt (quadratic) | 0.106 | 0.094 | 1.12 | .262 |
| Unskilled (vs. Skilled[a]) | 0.039 | 0.127 | 0.31 | .761 |
| Attempt (linear) × Unskilled | −0.005 | 0.115 | −0.04 | .967 |
| Attempt (quadratic) × Unskilled | −0.097 | 0.111 | −0.87 | .385 |

*Note.* "Attempt" denotes attempt relative to disengagement (−3, −2, −1), coded as an ordered factor with orthogonal polynomial contrasts. [a] Reference category.
